# Mechanisms and functions of respiration-driven gamma oscillations in the primary olfactory cortex

**DOI:** 10.1101/2022.04.24.489324

**Authors:** Joaquín González, Pablo Torterolo, Adriano BL Tort

**Affiliations:** Departamento de Fisiología, Facultad de Medicina, Universidad de la República, Montevideo, 11200, Uruguay; Brain Institute, Federal University of Rio Grande do Norte, Natal, RN 59056, Brazil

**Keywords:** neural oscillations, breathing, cell assemblies, sparse coding, optogenetics, winner-take-all

## Abstract

Gamma oscillations are believed to underlie cognitive processes by shaping the formation of transient neuronal partnerships on a millisecond scale. These oscillations are coupled to the phase of breathing cycles in several brain areas, possibly reflecting local computations driven by sensory inputs sampled at each breath. Here, we investigated the mechanisms and functions of gamma oscillations in the piriform (olfactory) cortex of awake mice to understand their dependence on breathing and how they relate to local spiking activity. Mechanistically, we find that respiration drives gamma oscillations in the piriform cortex, which correlate with local feedback inhibition and result from recurrent connections between local excitatory and inhibitory neuronal populations. Moreover, respiration-driven gamma oscillations are triggered by the activation of mitral/tufted cells in the olfactory bulb and are abolished during ketamine/xylazine anesthesia. Functionally, we demonstrate that they locally segregate neuronal assemblies through a winner-take-all computation leading to sparse odor coding during each breathing cycle. Our results shed new light on the mechanisms of gamma oscillations, bridging computation, cognition and physiology.

## Introduction

Since the pioneer works of Adrian in 1942 and Freeman in 1980, gamma oscillations have been one of the most studied brain rhythms (Bastos et al., 2020, 2015; Bragin et al., 1995; Buzsáki and Wang, 2012; Csicsvari et al., 2003; Fries et al., 2007; Gray et al., 1989; Sirota et al., 2008; Vinck et al., 2010; Womelsdorf et al., 2012, 2006). Gamma is believed to be critical for a variety of cognitive functions such as sensory processing (Fries et al., 2001), memory (Fernández-Ruiz et al., 2021), navigation (Colgin et al., 2009), and conscious awareness (Rodriguez et al., 1999). At the cellular scale, gamma rhythms modulate spiking activity, shaping the formation of transient neuronal partnerships, the so-called cell assemblies (Buzsáki, 2010). Computational models show that gamma depends on local inhibitory-inhibitory or excitatory-inhibitory interactions (Tort et al., 2007; Wang and Rinzel, 1992) and ultimately emerges from synchronous inhibitory postsynaptic potentials (Buzsáki and Wang, 2012). However, despite the initial evidence for their underlying principles being general, gamma oscillations are not a monolithic entity but actually encompass a diversity of rhythms observed experimentally (Lopes-Dos-Santos et al., 2018; Scheffer-Teixeira et al., 2012; Schomburg et al., 2014; Zhong et al., 2017). This warrants studying the mechanisms and functions of these oscillations in specific brain regions *in vivo*.

Gamma oscillations usually appear nested within slower rhythms, a phenomenon known as cross-frequency coupling, in which the amplitude of gamma waxes and wanes depending on the phase of a slow oscillation (Canolty and Knight, 2010; Lisman and Jensen, 2013). Important examples are the coupling of specific gamma sub-bands to the hippocampal theta rhythm (Cavelli et al., 2020; Sirota et al., 2008; Tort et al., 2009, 2008) or to the phase of breathing cycles (Cavelli et al., 2018; Ito et al., 2014; Zhong et al., 2017). Regarding the latter, it is worth noting that respiration-entrained brain rhythms depend on nasal airflow and are not a consequence of the respiratory pattern generation in the brainstem (Lockmann et al., 2016; Moberly et al., 2018; Yanovsky et al., 2014). Therefore, it seems likely that respiration-entrained gamma activity arises from local computations driven by sensory inputs sampled at each breath and thus plays a major role in cognition.

A promising area to study this hypothesis is the piriform cortex (PCx), which constitutes the primary olfactory area in the rodent brain (Bolding and Franks, 2018; Stettler and Axel, 2009) and exhibits prominent gamma oscillations (Bressler and Freeman, 1980; Courtiol et al., 2019; Freeman, 1960; Freeman and Skarda, 1985; Kay et al., 2009; Kay and Freeman, 1998; Litaudon et al., 2008; Mori et al., 2013; Vanderwolf, 2000). Most of our understanding of piriform oscillations comes from the studies of Walter Freeman in the 20th century (Barrie et al., 1996; Eeckman and Freeman, 1990; Freeman, 1968, 1960, 1959). Freeman characterized gamma activity in terms of its topography, frequency range, and relationship to unitary activity and behavior. These observations led him to hypothesize that these oscillations constitute a fundamental sensory processing component that emerges from an excitatory-inhibitory feedback loop. However, despite his influential insights, Freeman’s conjectures could not be conclusively tested due to the technological limitations of his time. Thus, we still lack compelling experimental demonstrations of how gamma generation depends on the interactions between the different piriform neuronal subpopulations or how gamma relates to odor representations encoded as cell assemblies. Therefore, the mechanisms of gamma oscillations in the PCx and their functional role in olfaction need to be studied under the lens of modern experimental and analytical tools (Courtiol et al., 2019; Kay et al., 2009; Mori et al., 2013).

In this report, we study gamma oscillations in the PCx of the awake mouse. We took advantage of modern genetic tools, which accurately identify neuronal populations and precisely modify the local connectivity of the PCx, thus enabling an unprecedented study of the mechanisms and functions of its oscillatory activity. We found that respiration drives gamma oscillations in this region, which derive from feedback inhibition and depend on recurrent connections between local excitatory and feedback inhibitory populations. This loop is triggered by the projection of mitral/tufted cells in the olfactory bulb onto principal cells of the PCx. As functional consequences, we show that respiration-driven gamma oscillations determine odor-assembly representations through a winner-take-all computation taking place within breathing cycles.

## Results

### Respiration drives gamma oscillations in the piriform cortex

To understand the mechanisms and functions of gamma oscillations and their relationship with respiration in the mouse brain, we analyzed local field potentials (LFP) from the PCx recorded simultaneously with the respiration signal (Fig 1A). The dataset was collected by Bolding and Franks (2018) and generously made available through CRCNS (http://crcns.org, pcx-1 dataset). Figure 1B depicts the LFP and respiration signal from a representative animal; notice that low-gamma oscillations (30-60 Hz) emerge following inhalation start. These low-gamma oscillations, already evident in raw recordings, are part of a larger gamma peak (30-100 Hz) in the LFP power spectrum (Fig 1C), reflecting a true oscillation (Yuval-Greenberg et al., 2008). Consistent with Figure 1B, only the low-gamma sub-band couples to the respiration cycle across animals, as its amplitude is modulated by both a 2-3 Hz LFP rhythm coherent to respiration (Fig 1D top panel; see also Figs S1 and S2) and by respiration itself (Fig 1D bottom panel). Thus, respiration entrains low-gamma oscillations in the PCx.

**Fig 1.**
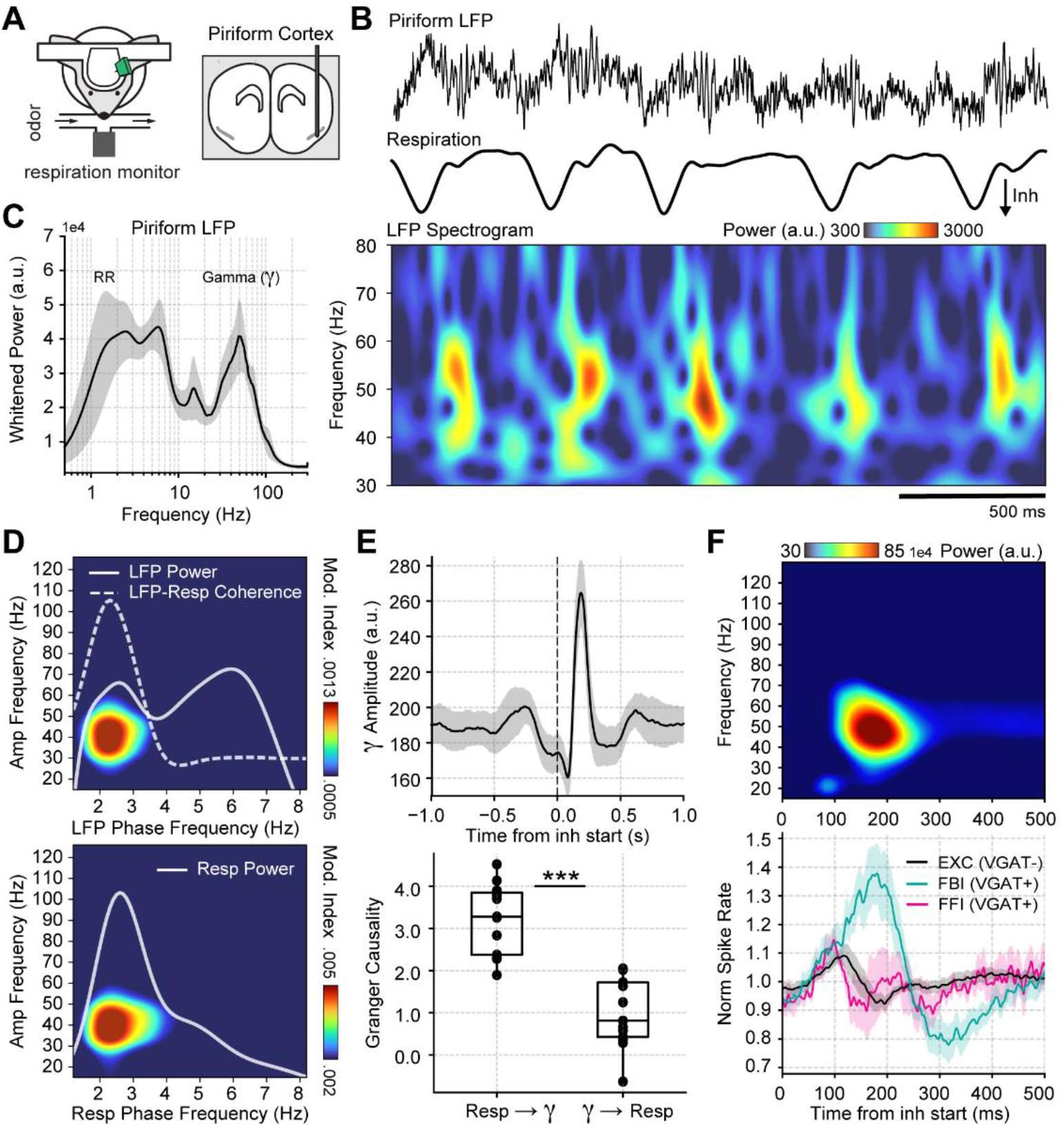
Respiration drives feedback inhibition-based gamma oscillations in the piriform cortex. **A** Experimental scheme and probe localization (modified from Bolding and Franks, 2018). **B** Example of simultaneously recorded LFP (top) and respiration (middle) signals, along with the LFP wavelet spectrogram (bottom). Notice prominent rhythmical appearance of gamma oscillations. **C** Average LFP power spectrum (± 2*SEM; n = 13 recording sessions from 12 mice). The spectrum was whitened by multiplying each value by the associated frequency. **D** Average phase-amplitude comodulogram using either the LFP (top) or the respiration (Resp; bottom) phase. Superimposed white lines show the LFP or Resp power spectrum (solid) and the LFP-Resp coherence (dashed). **E** Directionality analyses between Resp and the gamma envelope (30-60 Hz). Shown are the average (± SEM, n = 13 recording sessions from 12 mice) gamma envelope triggered by inhalation start (top), and the Granger causality for the Resp→gamma and gamma→Resp directions (bottom; boxplots show the median, 1st, 3rd quartiles, and the distribution range; each dot shows an individual mouse). **F** Respiration-evoked LFP responses. Top: average inhalation-triggered whitened spectrogram (n = 13 recording sessions from 12 mice). Bottom: Normalized spike rate (mean ± SEM, n = 15 recording sessions from 9 mice) of excitatory (EXC; VGAT-), feedback inhibitory (FBI; VGAT+) and feedforward inhibitory (FFI; VGAT+) neuronal populations triggered by inhalation.

To further characterize the interaction between respiration and low-gamma oscillations in the PCx, we performed directionality analyses (Fig 1E and Fig S3). The gamma amplitude envelope showed a peak ∼200 ms following the inhalation start, coinciding with a large positive cross-correlation peak between these signals, which suggests that respiration causes gamma. Consistent with these results, time-domain Granger causality was significantly higher in the respiration→gamma direction than in the opposite one (t(12) = 9.28, p<10^−6^). Together, these results show that respiration drives low-gamma oscillations in the PCx.

We analyzed the contribution of the different PCx neuronal populations to network low-gamma oscillations by first classifying single units according to the expression of a light-sensitive channelrhodopsin coupled to the vesicular GABA transporter (VGAT). This allowed to discriminate between VGAT-principal cells and VGAT+ inhibitory interneurons. Additionally, VGAT+ neurons were further classified into feedback inhibitory interneurons (FBI) and feedforward interneurons (FFI) according to their location relative to the principal cell layer (FBI tend to be located at a more superficial level within the cell layer 2/3; Bolding and Franks, 2018). Upon averaging the activity of each neuronal subpopulation, we found that the time-course of FBI firing rate changes correlates with the amplitude of low-gamma oscillations in both time and respiratory phase (Fig 1F and Fig S3). In contrast, principal cells and FFI spike earlier within the respiratory cycle and return to baseline during the gamma amplitude peak (Fig 1F and Fig S3). Notice further that the gamma peak coincides with principal cell inhibition, as evidenced by their firing rate decrease. Therefore, we conclude that respiration-driven low-gamma oscillations in the PCx arise from feedback inhibition.

### Respiration-driven gamma oscillations depend on recurrent connections within the piriform cortex

We next studied the circuit mechanisms responsible for the respiration-driven gamma oscillations in the PCx. To that end, we analyzed PCx LFPs following the selective expression of the tetanus toxin light chain in principal cells of a targeted hemisphere (TeLC ipsi; Fig 2A). Under this approach, TeLC expression blocks excitatory synaptic transmission without affecting cellular excitability (Bolding and Franks, 2018), allowing us to study the local computations underlying the respiration-driven gamma. Figure 2B shows that TeLC ipsi LFPs had a large significant reduction in low-gamma oscillations with respect to either control mice (t(19) = 4.59, p<0.0001) or the contralateral PCx (TeLC contra, not infected; t(12) = 3.79, p = 0.0012). Moreover, TeLC ipsi LFPs also showed a significant decrease in respiration-gamma coupling compared to control (t(19) = 10.52, p<10^−8^) or TeLC contra LFPs (t(12) = 2.78, p = 0.008) (Fig 2C,D), despite the respiration signal still reaching the PCx (note the dotted white traces in Fig 2C showing LFP-Resp coherence). These results demonstrate that gamma oscillations and their coupling to respiration depend on local recurrent excitatory connections within the PCx. Noteworthy, respiration-driven low-gamma oscillations also depend on the cognitive state since ketamine/xylazine anesthesia abolishes them (Fig S4).

**Fig 2.**
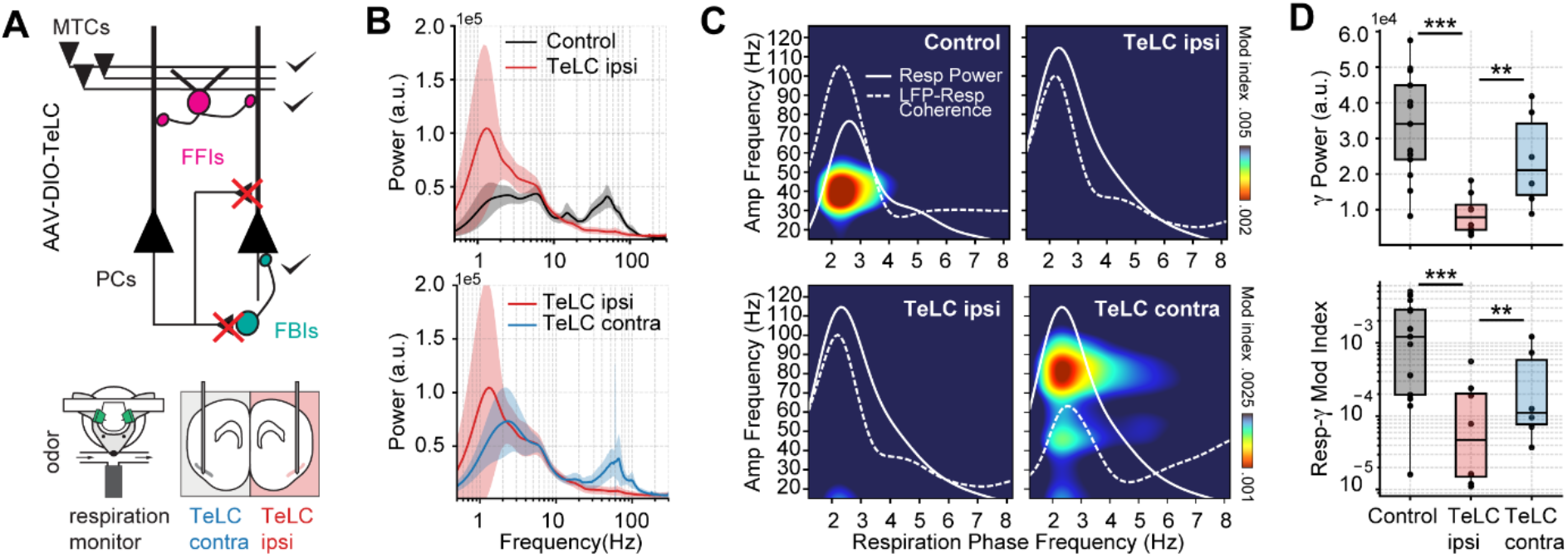
Respiration-driven gamma oscillations depend on recurrent connections within the piriform cortex. **A** Schematic of circuit changes after TeLC expression in principal cells (PCs) of the piriform circuit (MTCs: mitral cells; FFIs: feedforward interneurons; FBIs: feedback interneurons). Recordings were made both ipsi- and contralaterally to the TeLC expression (modified from Bolding and Franks, 2018). **B** Average (± 2*SEM) power spectra for control and TeLC-infected animals (Control, n = 13 recording sessions from 12 mice; TeLC ipsi, n = 8 recording sessions from 8 mice; TeLC contra, n = 6 recording sessions from 6 mice). Notice that local TeLC expression abolishes ipsilateral gamma oscillations in the PCx. **C** Average respiration-LFP comodulograms for control and TeLC-infected animals. Respiration power and LFP-respiration coherence are shown superimposed (same scale across plots). **D** Boxplots showing gamma power (top) and the Resp-low gamma modulation index (bottom) for control and TeLC-infected animals.

### Olfactory bulb mitral-cell projections trigger feedback inhibition-based gamma oscillations in the piriform cortex

After confirming that respiration-driven gamma oscillations depend on recurrent connections formed by principal cells, we asked how PCx inputs affect gamma generation. We expected mitral/tufted cell activation in the olfactory bulb (OB) to trigger similar low-gamma oscillations since these projections convey the respiratory inputs to the PCx (Pashkovski et al., 2020). Consistently, optogenetic activation of the OB (Thy-Control) triggered piriform low-gamma oscillations, which matched the laser time-course (Fig 3A, top panel). Interestingly, TeLC ipsi LFPs showed almost no gamma activity following laser onset (Fig 3A, middle panel), while TeLC contra LFPs still exhibited gamma activity upon stimulation of the OB (Fig 3A, bottom panel). Analyzing the group response, we found a significant reduction of low-gamma activity following the laser onset in TeLC ipsi LFPs compared to control (t(17) = 4.98, p<0.0001) or TeLC contra LFPs (t(19) = 3.49, p = 0.0012) (Fig 3B,C), further confirming the importance of recurrent connections for gamma generation. Moreover, low-gamma power in control and TeLC contra LFPs increased with laser intensity, while it remained constant in TeLC ipsi LFPs (Fig 3D). These results thus experimentally prove two critical facts about respiration-driven piriform gamma oscillations. First, that respiration drives PCx low-gamma oscillations mediated by OB projections from mitral/tufted cells. Second, feedforward interneurons do not generate low-gamma, which requires the principal cells to excite local feedback interneurons.

**Fig 3.**
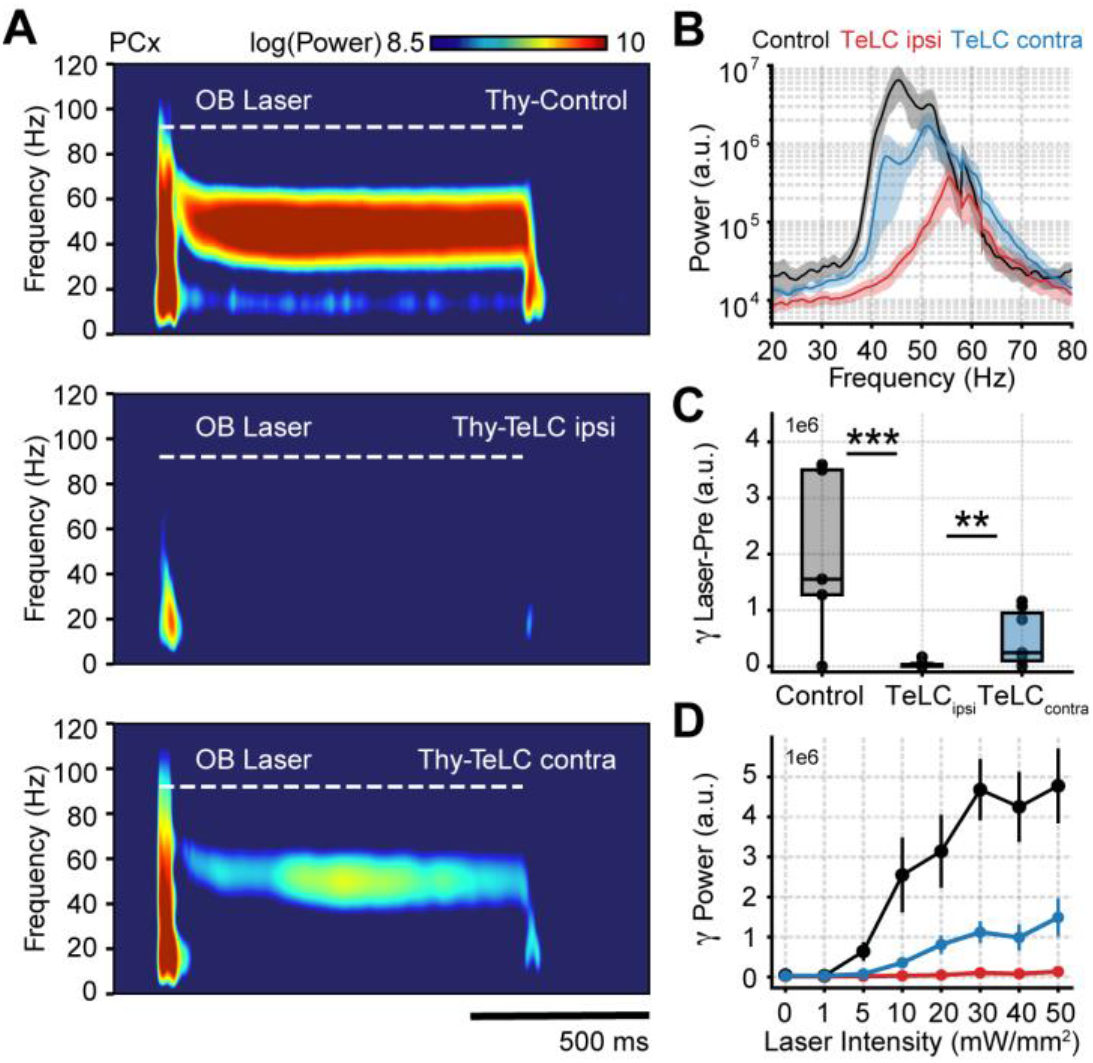
Piriform recurrent connections are necessary for OB mitral/tufted cells to trigger low-gamma oscillations. **A** Average PCx spectrograms during optogenetic stimulation of the olfactory bulb (OB). **B** Average gamma power during OB stimulation for the control (n = 5 recording sessions from 5 mice), TeLC ipsi (n = 14 recording sessions from 8 mice) and contralateral recordings (n = 7 recording sessions from 5 mice). Note that a logarithmic y-axis is employed here while subsequent plots use a linear scale. **C** Boxplots showing the gamma power difference between the laser and pre-laser periods. **D** Gamma power as a function of the laser intensity for each experimental condition (mean ± SEM).

### Piriform beta oscillations are evoked and do not require local computations

Having studied the mechanisms of respiration-driven low-gamma oscillations in awake mice, we next compared them with beta oscillations (10-30 Hz, Fig 4A,B), which have been widely associated with olfactory processing (Lepousez and Lledo, 2013; Martin et al., 2006; Poo and Isaacson, 2009). Beta oscillations occurred 100 ms before gamma onset, and, interestingly, showed a consistent phase resetting during each sniff cycle (Fig 4C top), resulting in a large amplitude envelope of the inhalation-trigged average of beta-filtered LFPs (Fig 4C middle), which is to say that beta was evoked at each cycle. The unfamiliar reader is referred to (Tallon-Baudry and Bertrand, 1999) for a discussion about evoked vs. induced oscillations. In short, oscillatory activity that shows up in the average filtered trace is said to be evoked since this requires phase consistency (or “resetting”) following each stimulus. On the other hand, oscillations that increase in amplitude following each stimulus (in our case, an inhalation), but that exhibit phase jitters from trial to trial, cannot be properly detected in the averaged trace due to peak-trough cancellations across trials. Such oscillations, referred to as “induced”, can only be detected by inspecting the average across all individual trial amplitudes (or spectrogram; see (Tallon-Baudry and Bertrand, 1999). This is the case of piriform low-gamma oscillations (compare Fig 4C middle and bottom panels) since they increase following each inhalation but exhibit time jitters in peak activity from cycle to cycle and no phase resetting (Fig 4C top). That is, our results show that gamma oscillations are not evoked but induced at each sniff cycle, contrasting therefore with the evoked beta oscillations.

**Fig 4.**
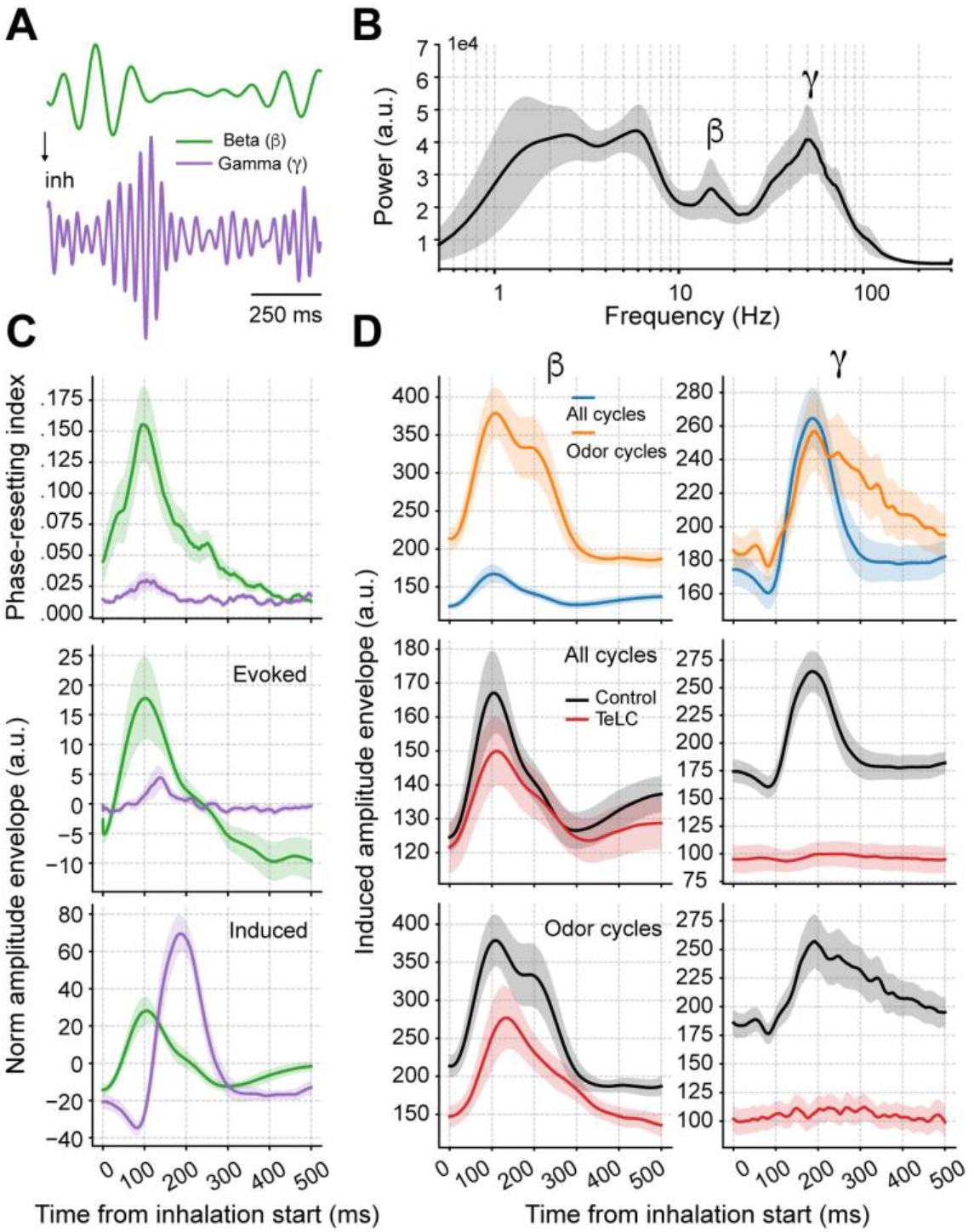
Piriform beta oscillations depend on odor delivery, are evoked at each cycle, and do not require recurrent local computations. **A** Filtered beta (10-20 Hz) and gamma (30-60 Hz) oscillations. **B** Average whitened LFP power spectrum (± 2*SEM; n = 13 recording sessions from 12 mice). **C** Top: Phase-resetting index for each oscillation. Middle: Normalized induced beta (green) and gamma (purple) amplitude triggered by inhalation. Bottom: Normalized evoked amplitude. The normalization consisted of removing the average amplitude across time. Traces show mean ± SEM. **D** Top: Average beta (left) and gamma (right) amplitude for all respiratory cycles (blue) and only for cycles with odor delivery (orange). Middle: Beta and gamma amplitude for all cycles under the control (black) or TeLC (red) condition. Bottom: As above, but for odor cycles. Shades represent the mean ± SEM. We note that because odorless cycles greatly outnumber odor ones (∼50-100 to 1), the oscillatory response to all-cycles and the odorless ones is virtually identical.

Next, we investigated beta and gamma responses to all sniffing cycles vs. the cycles in which a specific odorant was delivered (Fig 4D). Interestingly, odors increased the amplitude of beta oscillations but not gamma, though affected the duration of gamma activity at each sniffing cycle (further discussed below). Moreover, when comparing TeLC to control animals, we found that only gamma oscillations depended on the local piriform recurrent connections (as already shown in Fig 2), while beta oscillations were still present in infected animals (Fig 4D). In all, these results suggest that, in contrast to gamma, beta oscillations do not relate to local piriform computations and are likely of OB postsynaptic origin.

### Respiration-driven gamma oscillations determine odor-assembly representations through a winner-take-all computation

Finally, we investigated how respiration-driven gamma oscillations shape spiking patterns during odor sampling. First, we compared the average gamma responses of the odorless respiratory cycles against the cycles in which an odorant was presented. We found that odor delivery elicits longer-lasting gamma oscillations, irrespective of the specific odorant being delivered or its concentration (Fig S5), though with similar peak amplitude and peak latency within respiratory cycles (Fig 5A). Interestingly, odor delivery led to prolonged breaths; however, the longer respiratory cycle could not account for the prolonged gamma activity during odor sampling since no such effect was observed in odorless cycles matching odor breathing frequency (Fig S6). We then compared the firing rate of each piriform neuronal subtype during odorless and odor cycles (Fig 5B). Notably, feedback interneurons showed the most pronounced changes during odor cycles, substantially increasing their firing rate and prolonging their spiking period above baseline, thus mirroring the increase in gamma duration.

**Fig 5.**
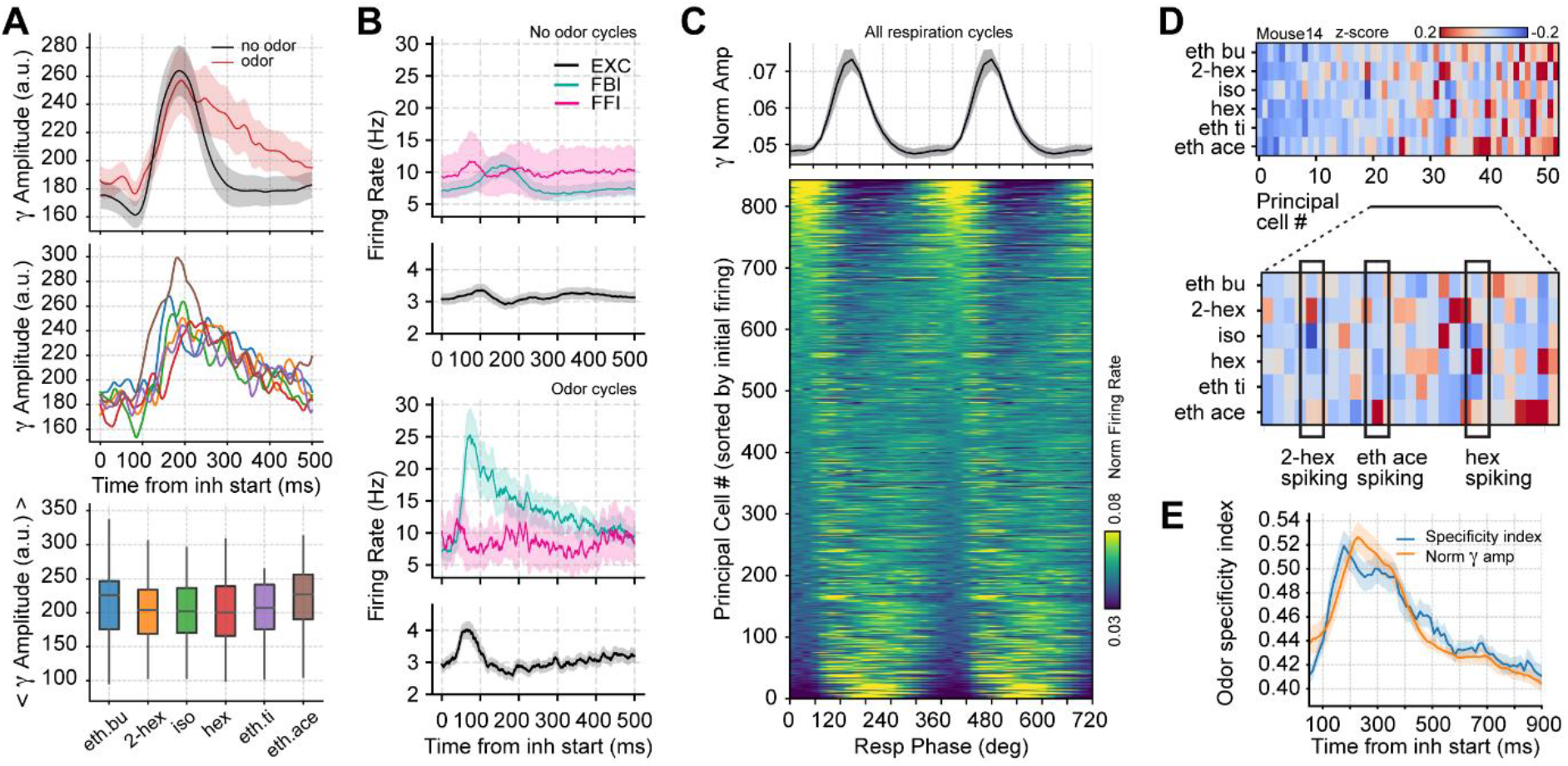
Respiration-driven gamma oscillations relate to single-cell spiking specificity to odors. **A** Top: Induced gamma amplitude triggered by the inhalation start. Shown are the average gamma response to odorless respiratory cycles and cycles containing odor sampling (n = 13 recording sessions from 12 mice). Middle: Average gamma amplitude for six different odorants. Bottom: Boxplots showing the mean gamma amplitude for the different odorants. **B** Neuronal firing rates (mean ± SEM, n = 15 recording sessions from 9 mice) during odorless (top) and odor cycles (bottom). **C** Principal cell spiking during each phase bin of the respiration cycle (bottom; 0 degree corresponds to the start of the inhalation); neurons are sorted according to the normalized firing rate in the first bin. Gamma power is shown on top (mean ± SEM, n = 15 recording sessions from 9 mice). **D** Z-scored firing rate at the gamma peak in response to different odors for a representative mouse. Columns show the firing rates of each principal cell. The bottom panel shows a zoom-in view of the differential spiking activity across odors. **E** Odor specificity index and normalized gamma amplitude following inhalation start (mean ± SEM, n = 15 recording sessions from 9 mice; gamma traces were rescaled to fit the plot).

Next, we studied how gamma oscillations influence principal cell spiking, pooling together all recorded principal cells and analyzing their spiking as a function of the respiration phase. We found that the respiration phase modulated principal cell spiking within breathing cycles (Fig 5C), though the preferred spiking phase differed across neurons. Interestingly, while a large proportion of cells were inhibited during the same respiration phase as the maximal gamma amplitude, some cells increased their spiking coincidently with the gamma peak. Importantly, the cells with increased spikes during gamma depended on odor context: when analyzing the population response to six different odorants (0.3% v./v. concentration), we found a noticeable cell-odor gamma spike specificity, that is, some cells spiked during gamma for a particular smell and not for the others (Fig 5D,E).

The gamma spiking specificity supports the conjecture that respiration-driven gamma oscillations could mediate odor assembly representations. To further study this possibility, we analyzed cell-assembly compositions for each odor by measuring the contribution of each principal cell to the first principal component (PC) of the population response (El-Gaby et al., 2021; Lopes-dos-Santos et al., 2013; Trouche et al., 2016). For each odor, we found that only a small fraction of neurons showed a strong positive contribution to the 1st PC, thereafter referred to as winner cells, while the vast majority showed low negative weights, referred to as loser cells (Fig 6A,B). Notably, the few winning neurons determining the 1st PC activity changed from odor to odor (Fig 6A); in other words, there was strong orthogonality in the 1st PC weight distribution across different odors. Consistent with this, the 1st PC weights were not significantly correlated between odors (p>0.016, corrected by multiple comparisons) (Fig 6C). Interestingly, the losing cells were significantly more phase-locked to the gamma phase (Fig 6D), consistent with them being actively inhibited during the oscillation. There were no differences between the preferred gamma phase for winners and losers (Fig 6E). Moreover, when analyzing the spiking time-course of winner and loser cells, we found that both groups tended to spike immediately following inhalation start; nevertheless, once feedback inhibition was recruited, loser cells spiked below their average while the winner cells continued spiking above baseline (Fig 6F). Notably, loser cell inhibition correlated with gamma activity, which suggests that sparse odor assembly representations emerge from a winner-take-all process mediated by inhibitory gamma oscillations.

**Fig 6.**
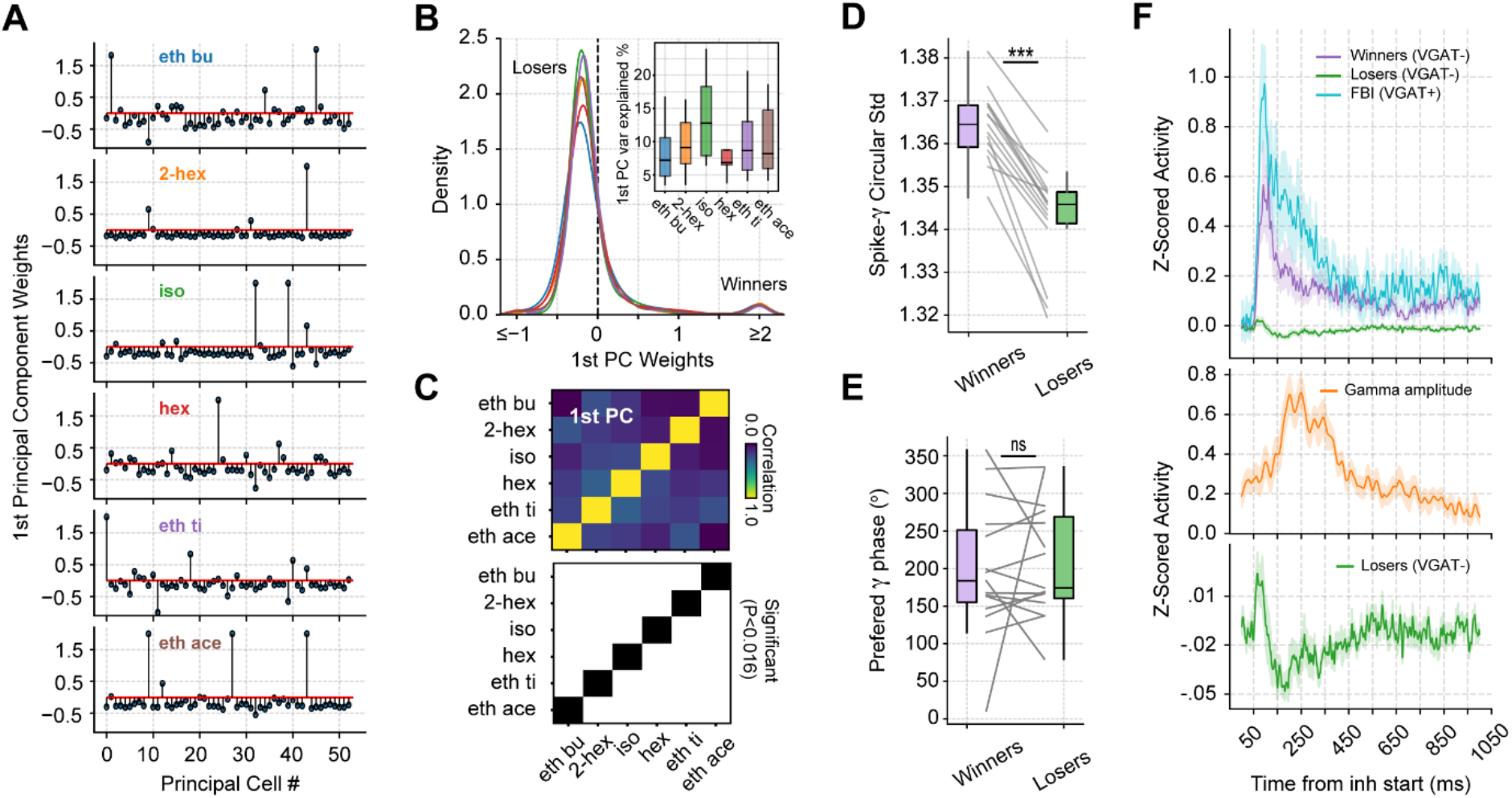
Gamma inhibition determines sparse odor-assembly representations through a winner-take-all computation. **A** Assembly weights for the 1st principal component (1st PC) in a representative mouse during the presentation of odorants. Notice different assembly compositions for the different odors. **B** Distribution of assembly weights for the 1st PC of each odor (all principal cell weights across sessions pooled together). Inset: % of variance explained by the 1st PC. **C** Correlation among 1st PC weights. The bottom panel shows the correlation statistics (thresholded at 0.016 according to the Bonferroni correction for multiple comparisons; black: statistically significant; white = non-significant). **D** Boxplots showing the circular standard deviation of the spiking gamma phase for winning and losing neurons. **E** Boxplots showing the preferred spiking gamma phase for winners and losers. **F** Average z-scored spiking activity for winners, losers, feedback inhibitory neurons (top), and average z-scored gamma amplitude envelope (middle). The bottom panel shows a y-axis zoom-in view of the spiking time-course of the loser neurons. N = 15 recording sessions from 9 mice.

Because a winner-take-all process selects a single odor representation while actively suppressing others, it effectively implements an XOR logic gate. As a consequence, we would expect odor decoding accuracy to increase during gamma inhibition. Following this reasoning, we first analyzed the correlation of population spiking vectors in response to different odorants and found that correlations significantly dropped during the gamma peak (Gamma vs. Pre: t(14) = −3.93, p = 0.0007; Gamma vs. Post: t(14) = −4.95, p = 0.0001; Fig 7A), confirming that odor representations diverge the most during gamma inhibition. We next tested odor decoding from the principal cell activity by training a supervised linear classifier using population spiking (i.e., spike vectors) at different respiratory cycle phases. Consistent with a winner-take-all mechanism, we found that variations in odor decoding accuracy matched the low-gamma activity time-course (Fig 7B,C); there was significant correlation between these variables (r = 0.47, p<10^−63^). Importantly, decoding accuracy significantly increased around the gamma peak compared to time windows before or after it (Gamma vs. Pre: t(14) = 5.20, p<0.0001; Gamma vs. Post: t(14) = 5.97, p<0.0001, Fig S7; see also Fig 7C). This result was robust among different classification algorithms (Fig S7), confirming that gamma inhibition generates a sparse PCx representation during odor processing. In all, these findings thus demonstrate that respiration-driven gamma oscillations provide an optimal temporal window for olfaction.

**Fig 7.**
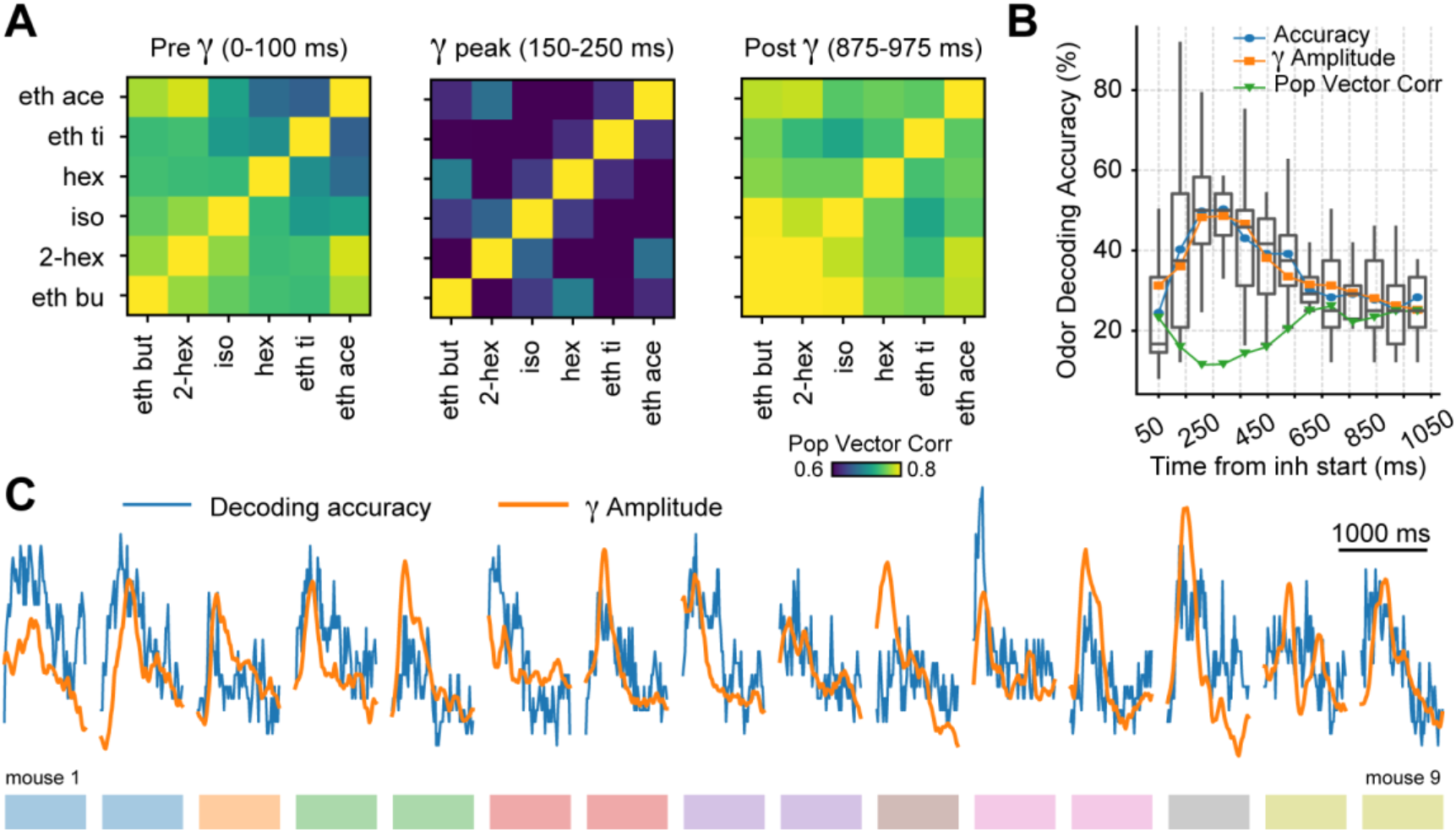
Gamma oscillations provide a privileged window for odor decoding. **A** Population vector correlations between odor responses during 100-ms time windows before (Pre), during (Gamma) and after (Post) the gamma peak. Only principal cells were employed for this analysis. **B** Boxplots showing odor decoding accuracy following inhalation start employing 100-ms time bins (n = 15 recording sessions from 9 mice). The mean accuracy across mice is shown by the blue line; for comparisons, the orange line shows the average gamma amplitude (arbitrary scale) and the green line shows the population vector correlations between principal cells (see Material and methods). **C** Odor decoding accuracy (blue) and gamma amplitude (orange) time-courses for each mouse. The colored rectangles underneath the traces show mouse identity.

## Discussion

The present report demonstrates the mechanisms and functions of gamma oscillations in the PCx of awake mice. We show that respiratory inputs to this cortex elicit large feedback inhibitory low-gamma oscillations that aid odor coding. Our work provides critical information for understanding gamma oscillations in the olfactory system. First, our results support the excitatory-inhibitory models of gamma generation (Buzsáki and Wang, 2012; Fisahn et al., 1998; Freeman, 1964), highlighting the role of excitatory neurons in initiating the gamma cycle by driving feedback interneuron spiking. Our results thus experimentally demonstrate several of Walter Freeman’s hypotheses regarding gamma generation (Bressler and Freeman, 1980; Eeckman and Freeman, 1990; Freeman, 1968), and reveal similarities with the mechanisms generating gamma in the OB (Kay et al., 2009; Lepousez and Lledo, 2013; Mori et al., 2013). Second, we demonstrate that the PCx low-gamma oscillations are the extracellular correlates of a winner-take-all process, thus confirming previous theoretical conjectures (de Almeida et al., 2009) and linking this brain oscillation to an algorithmic operation (an XOR logic gate). Third, the winner-take-all interpretation provides a direct relationship between gamma oscillations and cell assemblies. Under this scenario, the low-gamma feedback inhibition is critical for segregating cell assemblies and generating a sparse, orthogonal odor representation. Concomitantly, the assembly recruitment would depend on excitatory-excitatory interactions among winner cells occurring simultaneously during gamma activity.

An important feature of the respiration-driven low-gamma oscillations is that the winners triggering them change according to odor context (Fig 6A). Thus, the involvement of different cells and spiking dynamics likely makes the recruitment of feedback inhibition a non-homogenous process across respiration cycles, leading to small time jitters in peak gamma activity. As an empirical end result, the observed low-gamma oscillations are not phase-locked from cycle to cycle; in other words, PCx low-gamma power is induced, but not evoked (Tallon-Baudry and Bertrand, 1999), at each respiratory cycle. In contrast, beta oscillations were evoked by the respiratory input to the PCx, leading to a prominent phase resetting ∼100 ms after the inhalation start (Fig 4). Further consistent with these results, respiration-triggered beta oscillations did not disappear in the TeLC condition and correlated in time with excitatory spiking (c.f. Fig 4 and Fig 1). Interestingly, a previous report showed that odors evoke PCx beta oscillations under urethane anesthesia, which, as here, occurred before global inhibition (Poo and Isaacson, 2009). Based on this, a role for piriform beta in odor representations has been proposed; nevertheless, here we show that beta oscillations are likely inherited and do not require local computations in the PCx, i.e., they are not locally generated. In contrast, our results demonstrate that the piriform gamma does arise locally (and are not present during anesthesia), suggesting that odor coding ultimately depends on low-gamma oscillations in the main olfactory cortex of the mouse.

The study of the role of the PCx in olfaction has recently been boosted by selective genetic techniques, allowing researchers to examine how the individual cellular components relate to olfaction (Franks et al., 2011; Nagappan and Franks, 2021). The PCx, also known as the olfactory cortex, receives inputs from the OB and constructs a robust odor representation (Bolding et al., 2020). This representation can be inferred from the population activity and has two major features. On the one hand, it separates odor identity from its concentration (Bolding and Franks, 2018, 2017; Roland et al., 2017). This process depends on large-scale piriform inhibition (Bolding and Franks, 2018) and thus correlates with low-gamma oscillations. On the other hand, the representation groups odors into perceptual categories, not necessarily related to the physical structure of the odor molecules (Pashkovski et al., 2020). Surprisingly, these odor assembly representations are not fixed but change with time (Schoonover et al., 2021), which might be a particular property of unstructured cortices such as the piriform cortex (Stettler and Axel, 2009). In light of this evidence, spike-field synchronization might be an essential coding strategy, as neural oscillations might provide an internal signal in which assembly representations can be organized and structured. Thus, we hypothesize that low-gamma oscillations provide a critical time window for odor assembly representations to be formed and modified through winner-take-all computations.

The role of neuronal oscillations in shaping spiking patterns has been widely discussed (Buzsáki, 2010). Among other functions, oscillations are thought to provide syntactical blocks in which spiking activity can be parsed into meaningful words. A reader (receiver) area can then decode such neuronal representations by following these syntactic rules. In favor of this hypothesis, our decoding analysis shows that, for a reader neuron, optimal odor decoding occurs during the gamma peak within breathing cycles. Interestingly, and further consistent with these results, a recent study in humans showed that the accurate perception and identification of odors depend on piriform gamma oscillations (Yang et al., 2022). Moreover, the behavioral response to odors also seems to depend on piriform gamma, and its suppression induces depressive-like behaviors in mice (Li et al., 2022). Therefore, respiratory-triggered gamma oscillations might constitute a fundamental building block for neuronal communication in the olfactory system.

Several reports show that the PCx is the source of gamma activity for widespread brain regions, including the striatum (Carmichael et al., 2017) and the anterior limbic system (Carmichael et al., 2019). Although the authors largely attribute such oscillations to piriform volume conduction, recent evidence shows that olfactory gamma synchronizes distant regions as well (Li et al., 2022). Based on our results, we postulate that genuine respiration-entrained gamma oscillations in downstream regions are triggered by the output of the winning piriform assemblies, leading to new local gamma winner-take-all processes. This provides a new look into long-range gamma synchronization, emerging naturally if winners generating gamma in one area consistently trigger winners generating gamma in a downstream area. Such a mechanism closely agrees with the theory and experiments reported by Schneider et al. (2021), which suggest that oscillatory power and connectivity are main drivers underlying inter-areal coherence.

A central element to this directional view of interregional synchronization is gamma coupling to respiration, as the slow oscillation ensures a flow of information from the most primary olfactory areas (OB, PCx) to the higher limbic areas (prefrontal cortex, amygdala). In accordance, a recent report showed that respiration-related brain oscillations drive sparse assemblies in the prefrontal cortex (Folschweiller and Sauer, 2022). Notably, these authors also found that inhibitory recruitment by assembly members was key for assembly segregation, suggesting a potential role for gamma in that area. Hence, the directional view of synchronization across regions places a major role in cross-frequency interactions driving communication, highlighting the relevance of interregional synchrony modulation by slower rhythms, for instance, theta (González et al., 2020) or respiration itself (Cavelli et al., 2020). In light of our findings, we propose the following model to understand the relationship between spiking activity, neural oscillations, and odor coding in the piriform cortex: 1) Following each sniff, olfactory information is processed in the OB and is transmitted to the piriform circuit through mitral/tufted cell spiking. These synchronous postsynaptic potentials cause the field beta oscillation, whose amplitude depends on the afferent volley. 2) The principal cells that better encode the olfactory stimulus excite feedback interneurons. 3) Feedback interneurons inhibit competing principal cells, causing the field gamma oscillation. 4) This winner-take-all process segregates cell assemblies and dictates a sparse piriform odor representation. Note that because the piriform cortex normalized odorant concentration (Bolding and Franks, 2018), the amplitude of gamma oscillations does not change. Hence, gamma oscillations provide an optimal temporal window to decode odor.

## Material and methods

### Datasets

We analyzed recordings of the PCx generously made available by Bolding and Franks through the Collaborative Research in Computational Neuroscience data sharing website (http://crcns.org, pcx-1 dataset). Detailed descriptions of the experimental procedures can be found in previous publications (Bolding et al., 2020; Bolding and Franks, 2018, 2017). All protocols were approved by the Institutional Animal Care and Use Committee of Duke University. Below we describe the analytical methods employed by us, and, for convenience, also the relevant experimental procedures from the original publications.

#### Animals

Adult mice were employed (>P60, 20-24 g), housed in single cages on a normal light-dark cycle. For the Cre-dependent TeLC group, offspring of Emx1-cre (+/+) breeding pairs were obtained from The Jackson Laboratory (005628). For the optogenetic experiments, the mice employed were: adult Thy1-ChR2-YFP (+/+), line 18 (Thy1-COP4/EYFP, Jackson Laboratory, 007612) and VGAT-ChR2-YFP (+/−), line 8 (Slc32a1-COP4*H134R/EYFP, Jackson Laboratory, 014548). For the combined optogenetics and TeLC expression experiments, adult offspring of Emx1-cre (+/+) mice crossed with Thy1-ChR2-YFP (+/+) mice were employed.

#### Adeno-associated viral vectors

For the TeLC experiments, AAV5-DIO-TeLC-GFP was expressed under CBA control (6/7 mice) or synapsin (1/7 mice), whose effects were similar and pooled together. Three 500 nL injections in the PCx (AP, ML, DV: +1.8, 2.7, 3.85; +0.5, 3.5, 3.8; −1.5, 3.9, 4.2; DV measured from brain surface) were employed to achieve TeLC expression. All recordings took place ∼14 days after the promoter injection. All viruses were obtained from the University of North Carolina-Chapel Hill (UNC Vector Core)

#### Data acquisition

The electrophysiological signals were recorded using 32-site polytrode acute probes (A1×32-Poly3-5mm-25s-177, Neuronexus) with an A32-OM32 adaptor (Neuronexus) through a Cereplex digital headstage (Blackrock Microsystems). For the optogenetic identification of GABAergic cells, a fiber-attached polytrode probe was employed (A1×32-Poly3-5mm-25s-177-OA32LP, Neuronexus). Data was acquired at 30 kHz, unfiltered, employing a Cerebus multichannel data acquisition system (BlackRock Microsystems). Respiration and experimental events were acquired at 2 kHz by analog inputs of the Cerebus system. The respiration signal was measured employing a microbridge mass airflow sensor (Honeywell AWM3300V), which was positioned opposite to the animal nose. Inhalation generated a negative airflow and thus negative changes in the voltage of the sensor output.

#### Electrode and optic fiber placement

A Patchstar Micromanipulator (Scientifica) was employed to position the recording probe in the anterior PCx (1.32 mm anterior and 3.8 mm lateral from bregma). Recordings were targeted 3.5–4 mm ventral from the brain surface at this position, and were further adjusted according to the LFP and spiking activity monitored online. The electrode sites spanned 275 µm along the dorsal-ventral axis. The probe was lowered until an intense spiking band was found, which covered 30–40% of electrode sites near the correct ventral coordinate, thus reflecting the piriform layer II. For the optogenetic experiments stimulating OB cells in Thy1-ChR2-YFP mice, the optic fiber was placed <500 µm above the OB dorsal surface.

#### Spike sorting and waveform characteristics

Spyking-Circus software was employed to isolate individual units (https://github.com/spyking-circus). All clusters which had more than 1% of ISIs violating the refractory period (< 2 ms) or appearing contaminated were manually removed. Units which showed both similar waveforms and coordinated refractory periods were merged into a single cluster. The unit position was characterized as the mean electrode position (across electrodes) weighted by the amplitude of the unit waveform on each electrode.

### Analytical Methods

For all analyses, we used Python 3 with numpy (https://numpy.org/), scipy (https://docs.scipy.org/), matplotlib (https://matplotlib.org/), sklearn (https://scikit-learn.org/stable/) and statsmodel (https://www.statsmodels.org/stable/index.html) libraries.

#### Preprocessing

Raw LFPs were decimated to a sampling rate of 2000 Hz to match the respiration signal sampling rate, employing the *decimate* scipy function. This function first low-pass filters the 30 kHz raw data and then downsamples it, avoiding aliasing. For each animal, we used the same channel (channel 17) for all analyses, though similar results are obtained for the rest of the channels given their high LFP redundancy (Fig S8).

#### Power spectrum

To analyze the LFP spectra, we computed Welch’s modified periodogram using *welch* scipy function. Specifically, we employed a 1-s moving window with half a window overlap, setting the numerical frequency resolution to 0.1 Hz (by setting the nfft parameter to 10 times the sampling rate). All spectra were whitened by multiplying each power value by its associated frequency, thus eliminating the 1/f trend. For Figure 1B, we computed the wavelet transform (*cwt* scipy function) of the LFP signal using a Morlet wavelet with 0.1 Hz resolution.

The triggered spectrograms were obtained first selecting all 500-ms LFP windows following inhalation start (time stamps provided in the dataset, see Bolding and Franks 2018 for further details) and then computing the spectrogram using the *spectrogram* scipy function. We used 100-ms windows with 90% overlap and a frequency resolution of 0.1 Hz. After computing each individual spectrogram, we whitened (i.e., multiplied by f) and averaged them to yield the mean spectrogram shown in Figures 1F and 3A.

#### Phase-amplitude coupling

We computed phase-amplitude comodulograms following the methods proposed by (Tort et al., 2010). We first band-pass filtered the LFP/respiration signal using the *eegfilt* function (Delorme and Makeig, 2004) adapted for Python 3 (available at github/joaqgonzar). We filtered between 20 to 130 Hz in 10-Hz steps to obtain the higher frequency components, and between 1 and 8 Hz in 1-Hz steps to obtain the slower frequency components. The phase (angle) and amplitude time series of the filtered signals were estimated from their analytical representation based on the Hilbert transform. We then binned the phase time-series into 18 bins and computed the mean amplitude of the fast signal for each bin. The amount of phase-amplitude coupling was estimated through the modulation index (MI) metric (Tort et al., 2010), MI = (Hmax-H)/Hmax, where Hmax is the maximum possible Shannon entropy for a given distribution (log(number of phase bins)) and H is the actual entropy of the amplitude distribution.

#### Individual cell spiking responses to inhalation

We convolved the spike times of each unit with a 10-ms Gaussian kernel (standard deviation of 15 ms) employing the *convolve* scipy function. For the normalized responses, the spiking response of a neuronal subpopulation (i.e., excitatory, FFI or FBI neurons) was normalized by dividing the firing rate by its average across time [phase] bins; in Figures 1F and S3, the results show the mean ± SEM across animals.

#### Directionality analysis

To study the directionality between the gamma envelope and respiration signal, we employed three different strategies: (1) we averaged the gamma envelope using the inhalation start as a trigger; (2) we computed the cross-correlogram, using the correlate scipy function, between the gamma envelope and the respiration signal; (3) we computed Granger Causality estimates, using the grangercausalitytest stats model function, with a 10-order VAR model and employing the log(F-stat) as the Granger Causality magnitude (Geweke, 1982).

#### Induced power, evoked power and phase-resetting index

To study induced power, we (1) filtered the recordings (beta: 10-20 Hz, low gamma: 30-60 Hz), (2) obtained their amplitude through the Hilbert transform, and (3) computed the inhalation-triggered average. To study evoked power, we (1) filtered the recordings, (2) obtained the inhalation-triggered average, and then (3) estimated the amplitude of the average signal through the Hilbert transform. For obtaining the phase-resetting index of the filtered signals, we used the same procedure as for computing the inter-trial coherence (Makeig et al., 2004) using 500-ms windows following inhalation start as a “trial”.

#### Odor specificity index

We defined the odor specificity index for single cell responses following (McNaughton et al., 1983) as SI = max(Fx)/(Fa+Fb+Fc+Fd+Fe+Ff), where F represents firing rate, a-f represent different odors, and max(Fx) is the firing rate for the odor that generates the largest response.

#### Principal component analysis

We employed the *PCA* sklearn function on concatenated time-series of the smoothed principal cell spiking activity using all 1000-ms windows following inhalation start during odor presentation. Each odor was treated separately. We analyzed only the first principal component, explaining the most variance, setting n_components = 1. Winners and losers were identified according to the weight each neuron exerted on the 1st principal component. Namely, neurons with a positive weight were labeled as winners and neurons with a negative weight as losers. For representation purpose (Fig 6B), large positive and negative weights were truncated at 2 and −1, respectively.

#### Gamma phase-locking and preferred spiking phase

To assess coupling between spikes and gamma phase, the circular standard deviation of the spiking gamma phases was calculated using the *scipy circstd* function. The average gamma spiking angle (circmean) was computed to obtain the preferred phase.

#### Population vector correlations

To infer the similarity/dissimilarity in the population response to the different odors, we analyzed the population vector correlations, defined as the Pearson correlation (*pearsonr* scipy function) measured using paired data from the same neurons to two odors. Correlations were computed between spiking vectors for each odor; the correlation time course shown in Figure 7B was obtained using 100-ms sliding windows (87.5% overlap) following inhalation start. For statistics, we averaged the correlation across odors and compared them among 3 time windows of interest (see Fig 7A): from 0-100 ms (Pre), from 150-250 ms (Gamma), and from 875-975 ms (Post).

#### Odor decoding

We employed a supervised linear classifier to decode the odor stimulus from the population spiking vectors. The classification algorithm was supplied with the average spiking of each principal cell in 100-ms sliding windows following the inhalation start. The classification algorithm was a linear support vector machine with a stochastic gradient descent optimization, implemented using the *sgdclassifier* sklearn function. Each mouse was trained and tested separately; for each time window, the training sample consisted of 2/3 of the data, and the test sample consisted of the remaining third. The division of training/test samples was randomized in order to avoid odor bias. For the statistical comparisons, we used the same 100-ms time windows employed in the population vector correlations (i.e., Pre, Gamma, and Post). To verify the robustness of the results, we also employed a linear discriminant analysis and a k-nearest neighbor classifier, which rendered similar results to the sgdclassifier algorithm (Fig S7).

#### Statistics

We show data as either mean ± SEM or regular boxplots showing the median, 1st, 3rd quartiles and the distribution range without outliers. We employed paired and unpaired t-test to compare between groups. We set p<0.05 to be considered significant (in the figure panels, *, ** and *** denote p<0.05, p<0.01 and p<0.001, respectively). For the population vector correlations as well as 1st PC weight correlations, we employed Pearson r correlation and corrected the statistical threshold through the Bonferroni method for multiple comparisons (p<0.016).

## Code and data availability

All the data employed is freely available at: http://crcns.org, pcx-1 dataset. All of the analysis routines are available upon request to the first author.

## Acknowledgments

We thank Kevin Bolding and Kevin Franks for making their data available and for insightful comments on our manuscript. We also thank Diego Laplagne for comments on the manuscript. JG was supported by Comision Academica de Posgrado (CAP), Programa de Desarrollo de Ciencias Básicas (PEDECIBA) and Comisión Sectorial de Investigación Científica (CSIC). PT was supported by PEDECIBA and CSIC. ABLT was supported by Conselho Nacional de Desenvolvimento Científico e Tecnológico (CNPq) and Coordenação de Aperfeiçoamento de Pessoal de Nível Superior (CAPES), Brazil.

## Author contributions

JG conceptualized the work, created and implemented all of the analysis routines and wrote the initial manuscript. PT and ABLT supervised all the stages of this work and suggested analysis directions. All authors wrote the final manuscript.

## Supplementary Material

**Fig S1.**
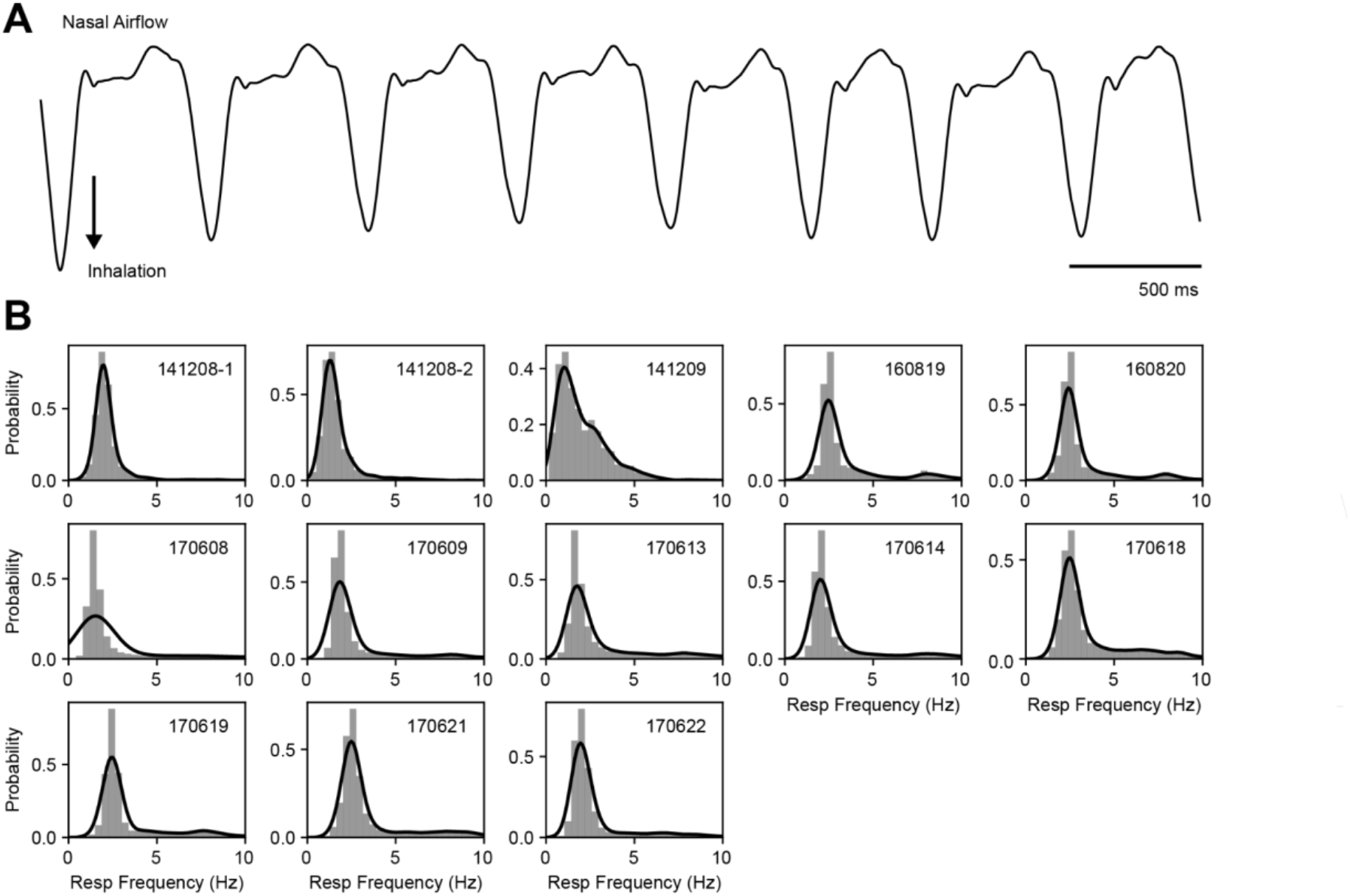
Similar respiratory frequency ranges across mice. **A** Example of a respiration recording measured through nasal airflow. **B** Respiratory frequency distribution for each mouse (insets show the mouse identification number). The instantaneous breathing frequency was estimated as the inverse of the inhalation period; bars depict probabilities for binned frequency intervals; solid lines show the kde distribution fit.

**Fig S2.**
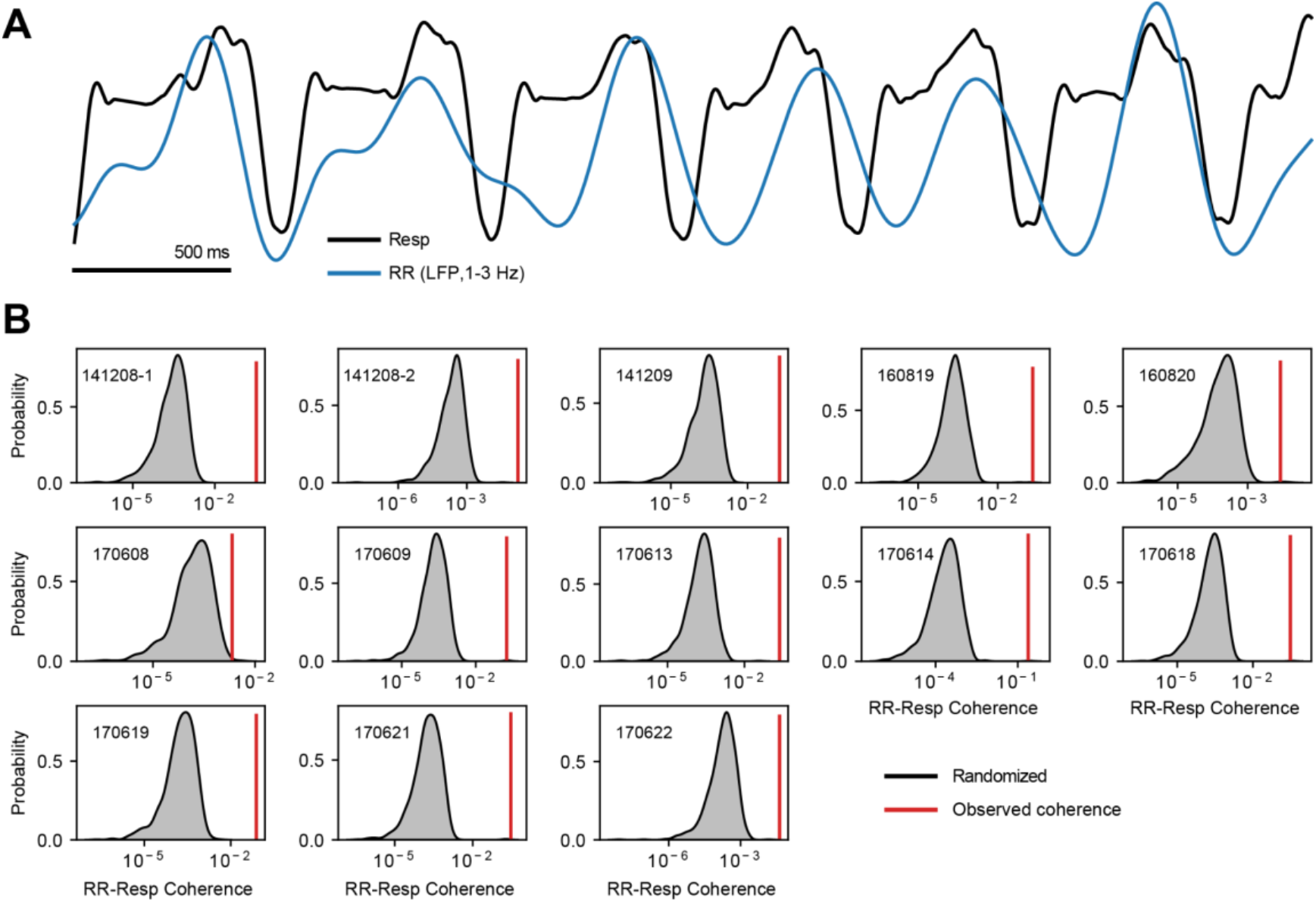
The 1-3 Hz LFP band is entrained by respiration. **A** Example of a respiration recording (Resp) along with the 1-3 Hz filtered LFP trace, which reflects the respiration-entrained rhythm (RR; see (Tort et al., 2018). **B** Surrogate RR-Resp coherence distributions (insets show the mouse identification number). Each distribution was obtained by computing the RR-Resp coherence using randomly circularly shifted respiration phases (1000 randomizations per mouse). The red vertical lines indicate the actual coherence values. All mice showed significant RR-Resp coherence (p < 0.01).

**Fig S3.**
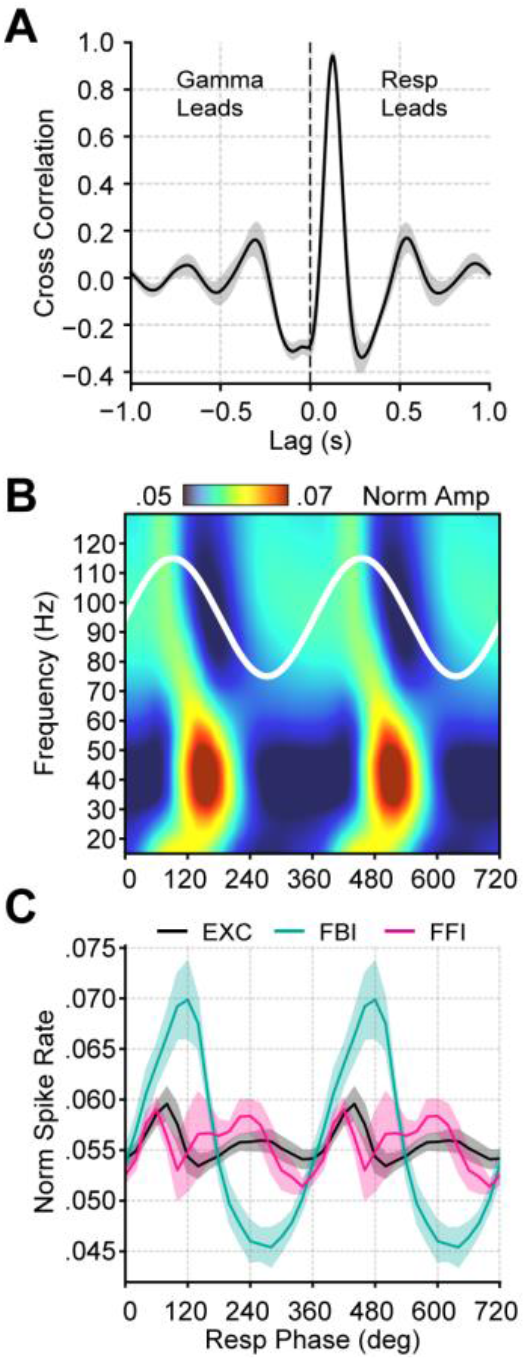
Respiration-entrained low-gamma oscillation in the piriform cortex. **A** Average (± SEM) cross-correlation between respiration (Resp) and the gamma envelope (n = 13 mice). **B** Normalized amplitude for LFP-filtered frequency components as a function of the Resp phase (average over 13 mice). **C** Spike rate (mean ± SEM) of excitatory (EXC; VGAT-) and feedback inhibitory (FBI; VGAT+) neuronal populations binned by the Resp phase (n = 15 mice).

## Supplementary Results Section

### Respiration-driven low-gamma oscillations fade under anesthesia

After unveiling the mechanisms responsible for the respiration-driven low-gamma activity in the PCx, we investigated whether these oscillations are state-dependent. To that end, we analyzed PCx LFP recordings during ketamine/xylazine anesthesia. Figure S4A shows the LFP spectrogram before and after ketamine/xylazine administration. Anesthesia promoted large-amplitude, slow LFP oscillations while greatly reducing gamma power, an effect that lasted ∼30 min (Fig S4A). These results can also be readily seen in the average power spectra of awake vs. anesthetized animals (Fig S4B), thus confirming that PCx low-gamma oscillations depend on the brain state. Next, we investigated if ketamine/xylazine anesthesia alters gamma coupling to respiration by analyzing respiration-LFP phase-amplitude comodulograms. We found that low-gamma oscillations are no longer coupled to respiration during general anesthesia (Fig S4C), despite the respiration-entrained rhythm being still present in the PCx LFP (Fig S4D,E). Intriguingly, the respiration-LFP comodulogram revealed coupling to a faster gamma activity (>80 Hz) under anesthesia (Fig S4C), which we deem likely to relate to the high-frequency oscillations evoked by sub-anesthetic doses of ketamine (Caixeta et al., 2013; Castro-Zaballa et al., 2018; Hunt et al., 2019), though it exhibited no power spectrum peak (c.f. Fig S4B).

**Fig S4.**
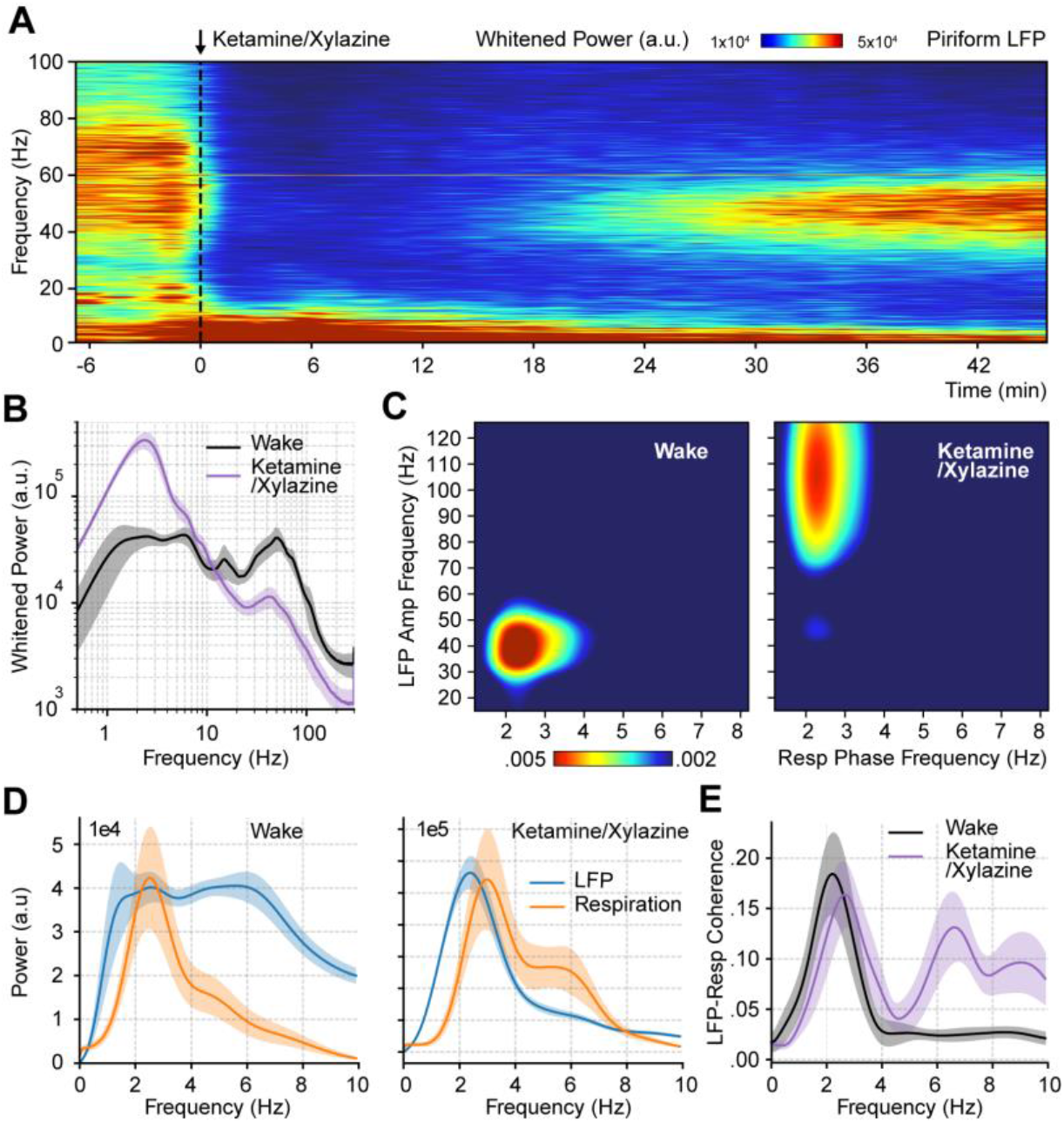
Ketamine/xylazine anesthesia abolishes low-gamma oscillations in the piriform cortex. **A** Spectrogram during ketamine/xylazine anesthesia in a representative animal (time of administration = 0 min). **B** Power spectra for awake and ketamine/xylazine-anesthetized animals (mean ± SEM; n = 11 mice). **C** Average respiration-LFP comodulogram for either brain state. **D** LFP (blue) and respiration (orange) power spectra (mean ± SEM) during wakefulness (left) and ketamine/xylazine anesthesia (right). **E** LFP-Resp coherence for each state (mean ± SEM; n = 10 mice).

**Fig S5.**
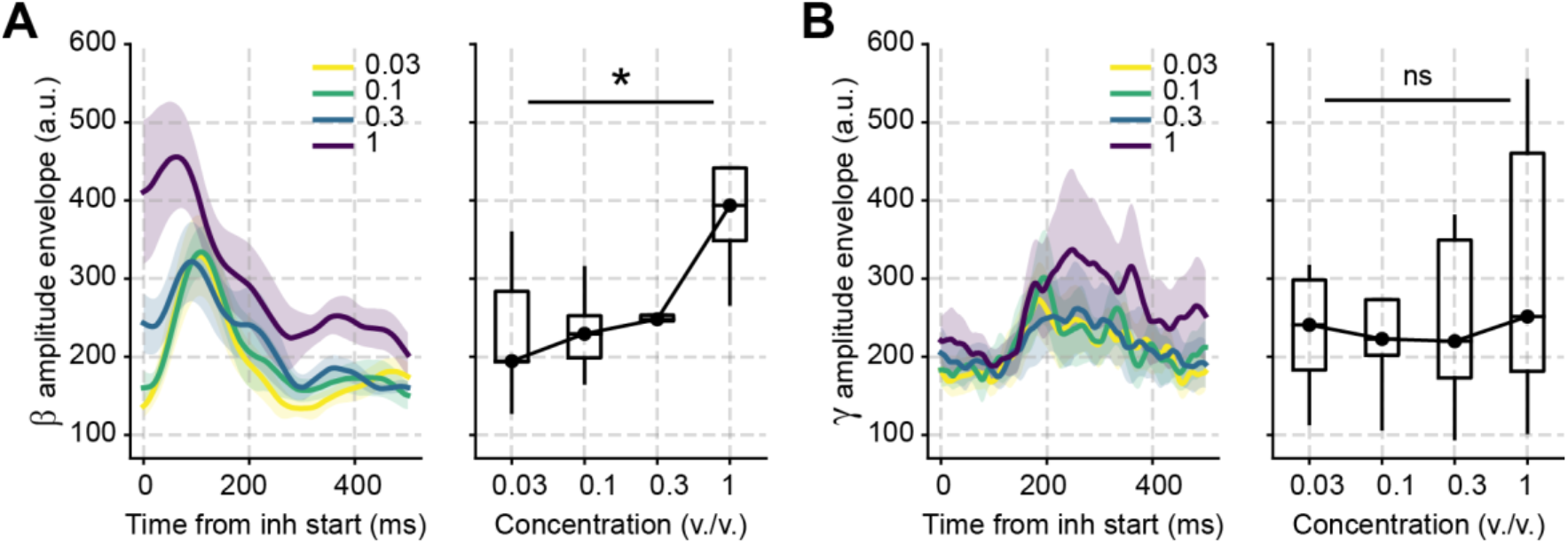
Beta amplitude depends on odor concentration. **A** Left: Average beta amplitude triggered by inhalation for ethyl butyrate and hexanal odors delivered at different concentrations (as labeled). Data from both odorants were averaged for these analyses. Right: Boxplots of the peak beta amplitude for each odor concentration (n = 5 mice). **B** As in A, but for gamma.

**Fig S6.**
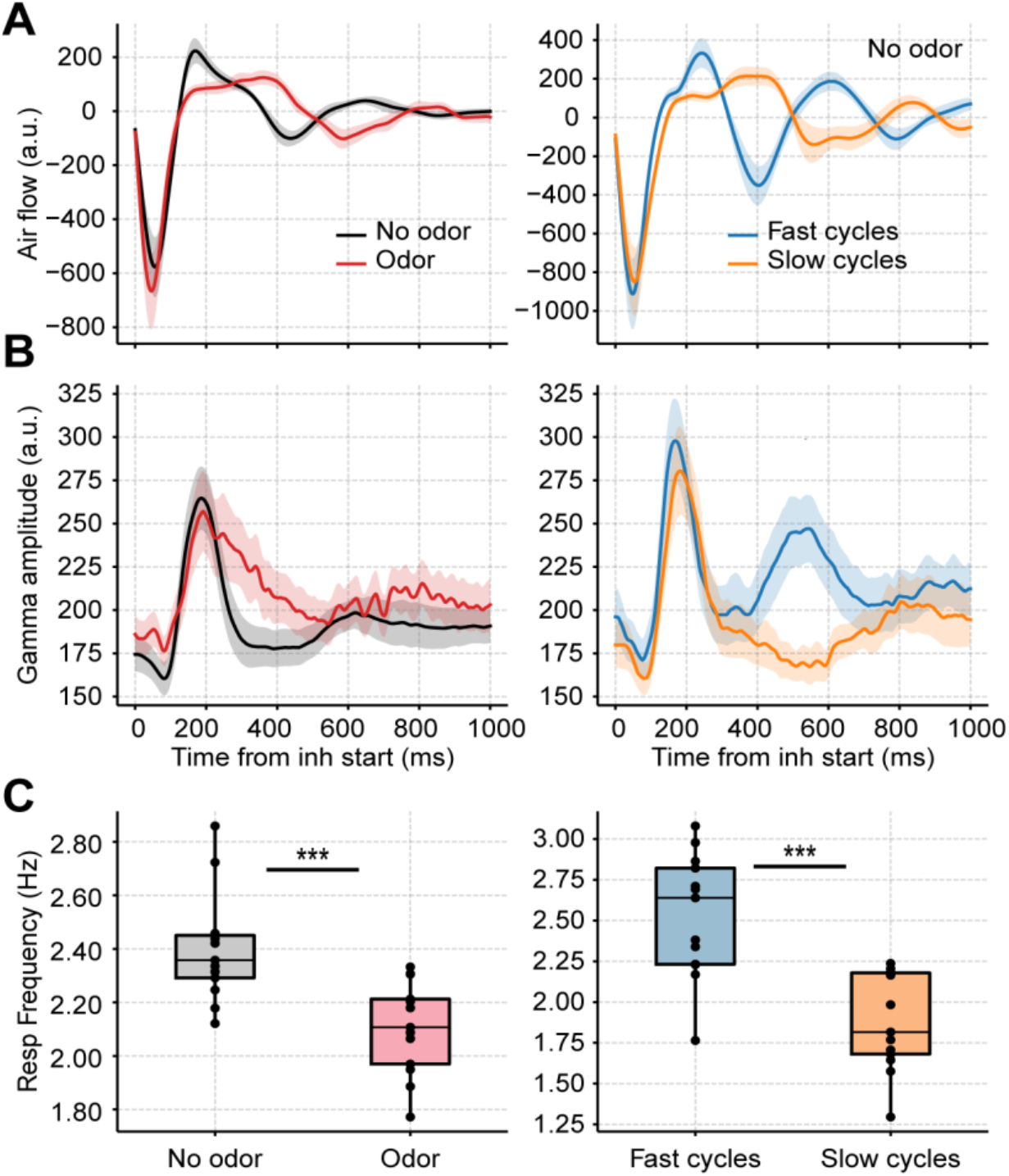
Longer breaths do not account for prolonged gamma activity in response to odors. **A** Average respiration waveforms triggered by the inhalation start (mean ± SEM). Left: Average breathing cycle in the presence (red) and absence (black) of odors. Right: Average over all fast (blue) and slow (orange) respiratory cycles in the absence of odors. Fast breaths were defined as those with an instantaneous frequency above the 75th percentile; slow breaths as cycles with an instantaneous frequency below the 25th percentile. **B** Average gamma amplitude triggered by inhalation start (mean ± SEM) for the same respiratory cycles as above. **C** Boxplots showing the average respiratory frequency during odor vs no odor cycles (left), and fast vs slow no odor cycles (right). Dots show individual animal values (n = 13 mice).

**Fig S7.**
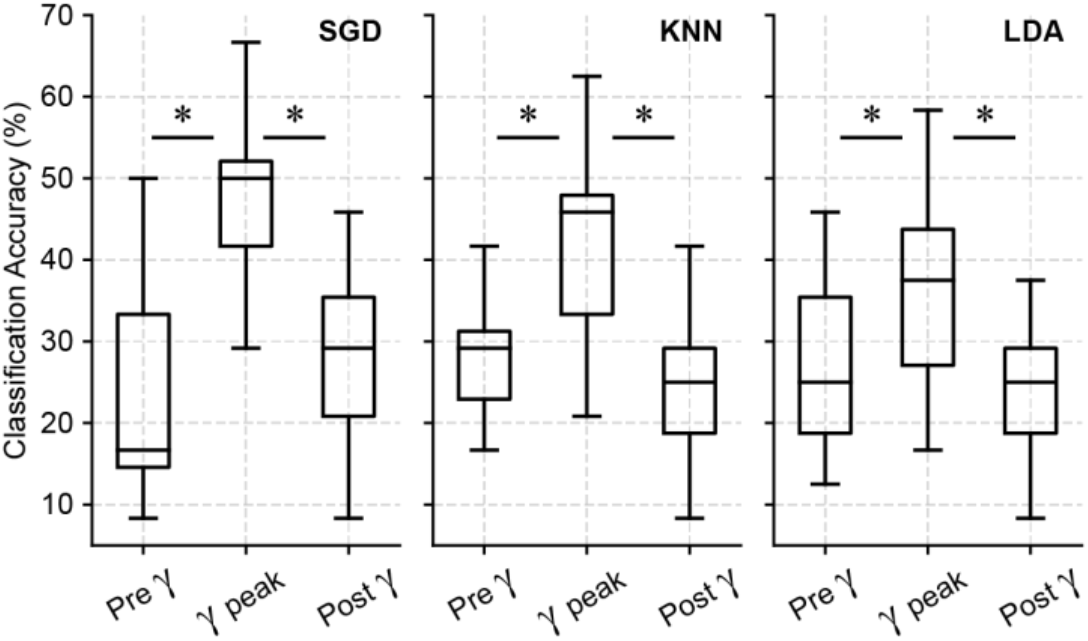
Odor decoding increases during the low-gamma peak. Boxplots showing odor decoding accuracy based on spiking activity measured using 100-ms windows before, during and after the respiration-driven gamma peak (see Material and methods). Each panel shows results from a different classification model (SGD: stochastic gradient descent classifier; KNN: k-nearest neighbors; LDA: linear discriminant analysis).

**Fig S8.**
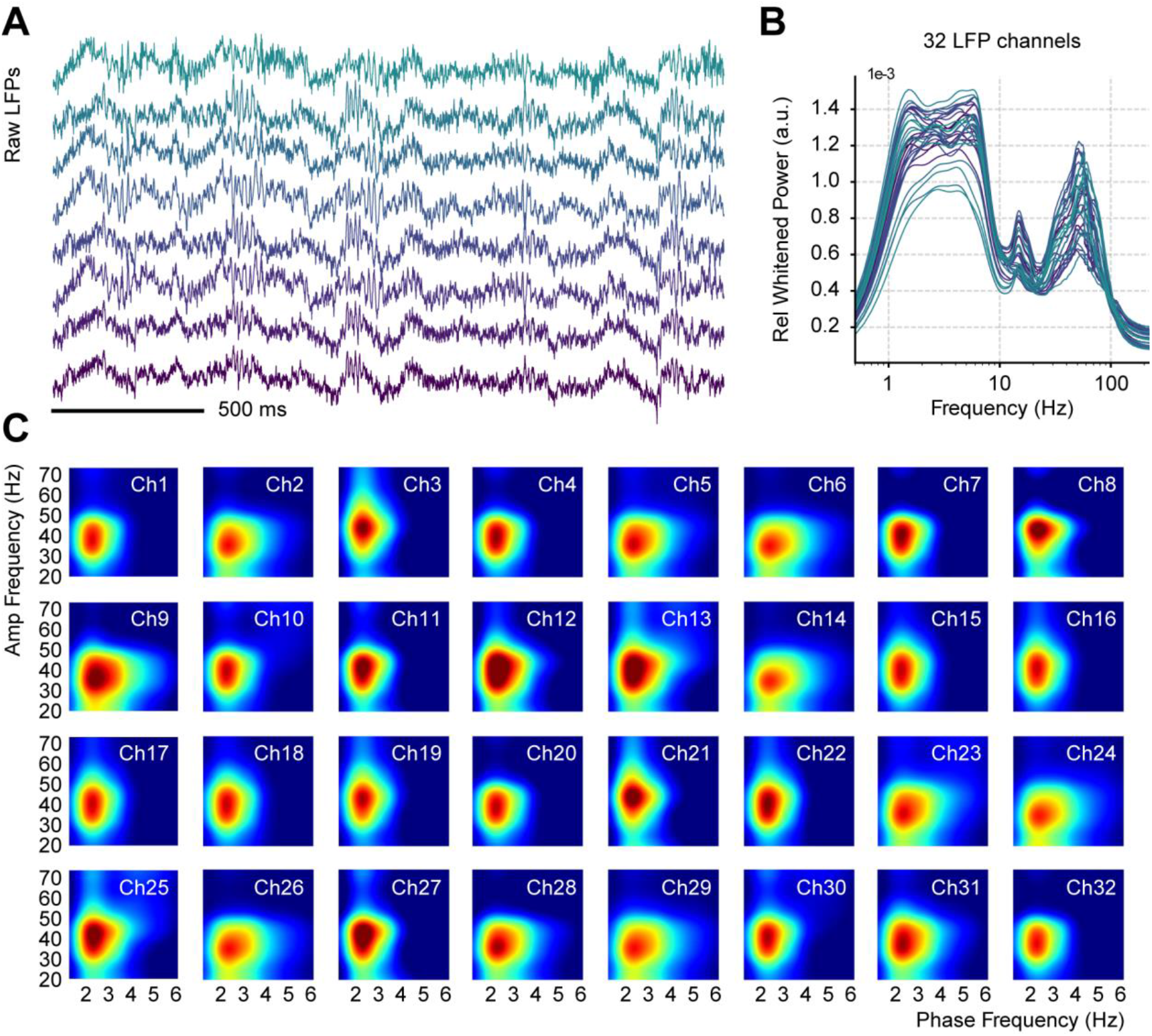
Respiration-driven gamma oscillations in the piriform cortex are evident across recording sites. **A** Simultaneous LFP recording examples from 8 channels. **B** Whitened normalized LFP power spectra for all 32 recording sites. The normalization consisted of dividing power values by the total power. Colors indicate the electrode number (from green to purple). **C** Phase-amplitude LFP comodulogram for each channel (individual color scales were optimized to depict RR-gamma coupling).

## Supplementary Methods

### Ketamine/xylazine anesthesia

Ketamine/xylazine (100/10 mg/kg) was administered intraperitoneally in 11 of the 13 mice from the control dataset, inducing stable anesthesia during a 30–45 min time period. A heating pad was used to maintain body temperature during this time. For all analyses, we employed the first 30 minutes following the slow-oscillation onset. A single animal was discarded since no electrophysiological markers of anesthesia were observed.

## References

Adrian ED. 1942. Olfactory reactions in the brain of the hedgehog. J Physiol 100:459–473.

Barrie JM, Freeman WJ, Lenhart MD. 1996. Spatiotemporal analysis of prepyriform, visual, auditory, and somesthetic surface EEGs in trained rabbits. J Neurophysiol 76:520–539.

Bastos AM, Lundqvist M, Waite AS, Kopell N, Miller EK. 2020. Layer and rhythm specificity for predictive routing. Proc Natl Acad Sci U S A 117:31459–31469.

Bastos AM, Vezoli J, Bosman CA, Schoffelen J-M, Oostenveld R, Dowdall JR, De Weerd P, Kennedy H, Fries P. 2015. Visual areas exert feedforward and feedback influences through distinct frequency channels. doi:10.1101/004804

Bolding KA, Franks KM. 2018. Recurrent cortical circuits implement concentration-invariant odor coding. Science 361. doi:10.1126/science.aat6904

Bolding KA, Franks KM. 2017. Complementary codes for odor identity and intensity in olfactory cortex. Elife 6. doi:10.7554/eLife.22630

Bolding KA, Nagappan S, Han B-X, Wang F, Franks KM. 2020. Recurrent circuitry is required to stabilize piriform cortex odor representations across brain states. Elife 9. doi:10.7554/eLife.53125

Bragin A, Jandó G, Nádasdy Z, Hetke J, Wise K, Buzsáki G. 1995. Gamma (40-100 Hz) oscillation in the hippocampus of the behaving rat. J Neurosci 15:47–60.

Bressler SL, Freeman WJ. 1980. Frequency analysis of olfactory system EEG in cat, rabbit, and rat. Electroencephalogr Clin Neurophysiol 50:19–24.

Buzsáki G. 2010. Neural syntax: cell assemblies, synapsembles, and readers. Neuron 68:362–385.

Buzsáki G, Wang X-J. 2012. Mechanisms of gamma oscillations. Annu Rev Neurosci 35:203–225.

Canolty RT, Knight RT. 2010. The functional role of cross-frequency coupling. Trends Cogn Sci 14:506–515.

Carmichael JE, Gmaz JM, van der Meer MAA. 2017. Gamma Oscillations in the Rat Ventral Striatum Originate in the Piriform Cortex. J Neurosci 37:7962–7974.

Carmichael JE, Yuen MM, van der Meer MAA. 2019. Piriform cortex provides a dominant gamma LFP oscillation in the anterior limbic system. bioRxiv. doi:10.1101/861021

Cavelli M, Castro-Zaballa S, Gonzalez J, Rojas-Líbano D, Rubido N, Velásquez N, Torterolo P. 2020. Nasal respiration entrains neocortical long-range gamma coherence during wakefulness. Eur J Neurosci 51:1463–1477.

Cavelli M, Rojas-Líbano D, Schwarzkopf N, Castro-Zaballa S, Gonzalez J, Mondino A, Santana N, Benedetto L, Falconi A, Torterolo P. 2018. Power and coherence of cortical high-frequency oscillations during wakefulness and sleep. Eur J Neurosci 48:2728–2737.

Colgin LL, Denninger T, Fyhn M, Hafting T, Bonnevie T, Jensen O, Moser M-B, Moser EI. 2009. Frequency of gamma oscillations routes flow of information in the hippocampus. Nature 462:353–357.

Courtiol E, Buonviso N, Litaudon P. 2019. Odorant features differentially modulate beta/gamma oscillatory patterns in anterior versus posterior piriform cortex. Neuroscience 409:26–34.

Csicsvari J, Jamieson B, Wise KD, Buzsáki G. 2003. Mechanisms of gamma oscillations in the hippocampus of the behaving rat. Neuron 37:311–322.

de Almeida L, Idiart M, Lisman JE. 2009. A second function of gamma frequency oscillations: an E%-max winner-take-all mechanism selects which cells fire. J Neurosci 29:7497–7503.

Delorme A, Makeig S. 2004. EEGLAB: an open source toolbox for analysis of single-trial EEG dynamics including independent component analysis. J Neurosci Methods 134:9–21.

Eeckman FH, Freeman WJ. 1990. Correlations between unit firing and EEG in the rat olfactory system. Brain Res 528:238–244.

El-Gaby M, Reeve HM, Lopes-Dos-Santos V, Campo-Urriza N, Perestenko PV, Morley A, Strickland LAM, Lukács IP, Paulsen O, Dupret D. 2021. An emergent neural coactivity code for dynamic memory. Nat Neurosci 24:694–704.

Fernández-Ruiz A, Oliva A, Soula M, Rocha-Almeida F, Nagy GA, Martin-Vazquez G, Buzsáki G. 2021. Gamma rhythm communication between entorhinal cortex and dentate gyrus neuronal assemblies. Science 372. doi:10.1126/science.abf3119

Fisahn A, Pike FG, Buhl EH, Paulsen O. 1998. Cholinergic induction of network oscillations at 40 Hz in the hippocampus in vitro. Nature 394:186–189.

Folschweiller S, Sauer J. 2022. Phase-specific pooling of sparse assembly activity by respiration-related brain oscillations. The Journal of Physiology. doi:10.1113/jp282631

Franks KM, Russo MJ, Sosulski DL, Mulligan AA, Siegelbaum SA, Axel R. 2011. Recurrent circuitry dynamically shapes the activation of piriform cortex. Neuron 72:49–56.

Freeman WJ. 1968. Relations between unit activity and evoked potentials in prepyriform cortex of cats. J Neurophysiol 31:337–348.

Freeman WJ. 1964. A linear distributed feedback model for prepyriform cortex. Exp Neurol 10:525–547.

Freeman WJ. 1960. Correlation of electrical activity of prepyriform cortex and behavior in cat. J Neurophysiol 23:111–131.

Freeman WJ. 1959. Distribution in time and space of prepyriform electrical activity. J Neurophysiol 22:644–665.

Freeman WJ, Skarda CA. 1985. Spatial EEG patterns, non-linear dynamics and perception: the neo-Sherringtonian view. Brain Res 357:147–175.

Fries P, Nikolić D, Singer W. 2007. The gamma cycle. Trends Neurosci 30:309–316.

Fries P, Reynolds JH, Rorie AE, Desimone R. 2001. Modulation of oscillatory neuronal synchronization by selective visual attention. Science 291:1560–1563.

Geweke J. 1982. Measurement of Linear Dependence and Feedback between Multiple Time Series. Journal of the American Statistical Association. doi:10.1080/01621459.1982.10477803

González J, Cavelli M, Mondino A, Rubido N, Tort AB, Torterolo P. 2020. Communication Through Coherence by Means of Cross-frequency Coupling. Neuroscience 449:157–164.

Gray CM, König P, Engel AK, Singer W. 1989. Oscillatory responses in cat visual cortex exhibit inter-columnar synchronization which reflects global stimulus properties. Nature 338:334–337.

Ito J, Roy S, Liu Y, Cao Y, Fletcher M, Lu L, Boughter JD, Grün S, Heck DH. 2014. Whisker barrel cortex delta oscillations and gamma power in the awake mouse are linked to respiration. Nat Commun 5:3572.

Kay LM, Beshel J, Brea J, Martin C, Rojas-Líbano D, Kopell N. 2009. Olfactory oscillations: the what, how and what for. Trends Neurosci 32:207–214.

Kay LM, Freeman WJ. 1998. Bidirectional processing in the olfactory-limbic axis during olfactory behavior. Behavioral Neuroscience. doi:10.1037/0735-7044.112.3.541

Lepousez G, Lledo P-M. 2013. Odor discrimination requires proper olfactory fast oscillations in awake mice. Neuron 80:1010–1024.

Li Q, Takeuchi Y, Wang J, Barcsai L, Pedraza LK, Kozák G, Nakai S, Kato S, Kobayashi K, Ohsawa M, Lőrincz ML, Devinsky O, Buzsaki G, Berényi A. 2022. Reinstating olfactory bulb derived limbic gamma oscillations alleviates depression. doi:10.1101/2022.02.01.478683

Lisman JE, Jensen O. 2013. The Theta-Gamma Neural Code. Neuron 77:1002–1016.

Litaudon P, Garcia S, Buonviso N. 2008. Strong coupling between pyramidal cell activity and network oscillations in the olfactory cortex. Neuroscience 156:781–787.

Lockmann ALV, Laplagne DA, Leão RN, Tort ABL. 2016. A Respiration-Coupled Rhythm in the Rat Hippocampus Independent of Theta and Slow Oscillations. J Neurosci 36:5338–5352.

Lopes-dos-Santos V, Ribeiro S, Tort ABL. 2013. Detecting cell assemblies in large neuronal populations. J Neurosci Methods 220:149–166.

Lopes-Dos-Santos V, van de Ven GM, Morley A, Trouche S, Campo-Urriza N, Dupret D. 2018. Parsing Hippocampal Theta Oscillations by Nested Spectral Components during Spatial Exploration and Memory-Guided Behavior. Neuron 100:940–952.e7.

Makeig S, Debener S, Onton J, Delorme A. 2004. Mining event-related brain dynamics. Trends Cogn Sci 8:204–210.

Martin C, Gervais R, Messaoudi B, Ravel N. 2006. Learning-induced oscillatory activities correlated to odour recognition: a network activity. Eur J Neurosci 23:1801–1810.

McNaughton BL, Barnes CA, O’Keefe J. 1983. The contributions of position, direction, and velocity to single unit activity in the hippocampus of freely-moving rats. Exp Brain Res 52:41–49.

Moberly AH, Schreck M, Bhattarai JP, Zweifel LS, Luo W, Ma M. 2018. Olfactory inputs modulate respiration-related rhythmic activity in the prefrontal cortex and freezing behavior. Nature Communications. doi:10.1038/s41467-018-03988-1

Mori K, Manabe H, Narikiyo K, Onisawa N. 2013. Olfactory consciousness and gamma oscillation couplings across the olfactory bulb, olfactory cortex, and orbitofrontal cortex. Front Psychol 4:743.

Nagappan S, Franks KM. 2021. Parallel processing by distinct classes of principal neurons in the olfactory cortex. Elife 10. doi:10.7554/eLife.73668

Pashkovski SL, Iurilli G, Brann D, Chicharro D, Drummey K, Franks KM, Panzeri S, Datta SR. 2020. Structure and flexibility in cortical representations of odour space. Nature 583:253–258.

Poo C, Isaacson JS. 2009. Odor Representations in Olfactory Cortex: “Sparse” Coding, Global Inhibition, and Oscillations. Neuron 62:850–861.

Rodriguez E, George N, Lachaux J-P, Martinerie J, Renault B, Varela FJ. 1999. Perception’s shadow: long-distance synchronization of human brain activity. Nature. doi:10.1038/17120

Roland B, Deneux T, Franks KM, Bathellier B, Fleischmann A. 2017. Odor identity coding by distributed ensembles of neurons in the mouse olfactory cortex. Elife 6. doi:10.7554/eLife.26337

Scheffer-Teixeira R, Belchior H, Caixeta FV, Souza BC, Ribeiro S, Tort ABL. 2012. Theta phase modulates multiple layer-specific oscillations in the CA1 region. Cereb Cortex 22:2404–2414.

Schneider M, Broggini AC, Dann B, Tzanou A, Uran C, Sheshadri S, Scherberger H, Vinck M. 2021. A mechanism for inter-areal coherence through communication based on connectivity and oscillatory power. Neuron 109:4050–4067.e12.

Schomburg EW, Fernández-Ruiz A, Mizuseki K, Berényi A, Anastassiou CA, Koch C, Buzsáki G. 2014. Theta phase segregation of input-specific gamma patterns in entorhinal-hippocampal networks. Neuron 84:470–485.

Schoonover CE, Ohashi SN, Axel R, Fink AJP. 2021. Representational drift in primary olfactory cortex. Nature 594:541–546.

Sirota A, Montgomery S, Fujisawa S, Isomura Y, Zugaro M, Buzsáki G. 2008. Entrainment of neocortical neurons and gamma oscillations by the hippocampal theta rhythm. Neuron 60:683–697.

Stettler DD, Axel R. 2009. Representations of Odor in the Piriform Cortex. Neuron. doi:10.1016/j.neuron.2009.10.008

Tallon-Baudry C, Bertrand O. 1999. Oscillatory gamma activity in humans and its role in object representation. Trends Cogn Sci 3:151–162.

Tort ABL, Komorowski R, Eichenbaum H, Kopell N. 2010. Measuring phase-amplitude coupling between neuronal oscillations of different frequencies. J Neurophysiol 104:1195–1210.

Tort ABL, Komorowski RW, Manns JR, Kopell NJ, Eichenbaum H. 2009. Theta-gamma coupling increases during the learning of item-context associations. Proc Natl Acad Sci U S A 106:20942–20947.

Tort ABL, Kramer MA, Thorn C, Gibson DJ, Kubota Y, Graybiel AM, Kopell NJ. 2008. Dynamic cross-frequency couplings of local field potential oscillations in rat striatum and hippocampus during performance of a T-maze task. Proc Natl Acad Sci U S A 105:20517–20522.

Tort ABL, Rotstein HG, Dugladze T, Gloveli T, Kopell NJ. 2007. On the formation of gamma-coherent cell assemblies by oriens lacunosum-moleculare interneurons in the hippocampus. Proc Natl Acad Sci U S A 104:13490–13495.

Trouche S, Perestenko PV, van de Ven GM, Bratley CT, McNamara CG, Campo-Urriza N, Black SL, Reijmers LG, Dupret D. 2016. Recoding a cocaine-place memory engram to a neutral engram in the hippocampus. Nat Neurosci 19:564–567.

Vanderwolf CH. 2000. What is the significance of gamma wave activity in the pyriform cortex? Brain Res 877:125–133.

Vinck M, Lima B, Womelsdorf T, Oostenveld R, Singer W, Neuenschwander S, Fries P. 2010. Gamma-phase shifting in awake monkey visual cortex. J Neurosci 30:1250–1257.

Wang X-J, Rinzel J. 1992. Alternating and synchronous rhythms in reciprocally inhibitory model neurons. Neural Comput 4:84–97.

Womelsdorf T, Fries P, Mitra PP, Desimone R. 2006. Gamma-band synchronization in visual cortex predicts speed of change detection. Nature 439:733–736.

Womelsdorf T, Lima B, Vinck M, Oostenveld R, Singer W, Neuenschwander S, Fries P. 2012. Orientation selectivity and noise correlation in awake monkey area V1 are modulated by the gamma cycle. Proc Natl Acad Sci U S A 109:4302–4307.

Yang Q, Zhou G, Noto T, Templer JW, Schuele SU, Rosenow JM, Lane G, Zelano C. 2022. Smell-induced gamma oscillations in human olfactory cortex are required for accurate perception of odor identity. PLoS Biol 20:e3001509.

Yanovsky Y, Ciatipis M, Draguhn A, Tort ABL, Brankačk J. 2014. Slow oscillations in the mouse hippocampus entrained by nasal respiration. J Neurosci 34:5949–5964.

Yuval-Greenberg S, Tomer O, Keren AS, Nelken I, Deouell LY. 2008. Transient induced gamma-band response in EEG as a manifestation of miniature saccades. Neuron 58:429–441.

Zhong W, Ciatipis M, Wolfenstetter T, Jessberger J, Müller C, Ponsel S, Yanovsky Y, Brankačk J, Tort ABL, Draguhn A. 2017. Selective entrainment of gamma subbands by different slow network oscillations. Proc Natl Acad Sci U S A 114:4519–4524.

## Supplementary References

Caixeta, F.V., Cornélio, A.M., Scheffer-Teixeira, R., Ribeiro, S., and Tort, A.B.L. (2013). Ketamine alters oscillatory coupling in the hippocampus. Sci. Rep. 3, 2348.

Castro-Zaballa, S., Cavelli, M.L., Gonzalez, J., Nardi, A.E., Machado, S., Scorza, C., and Torterolo, P. (2018). EEG 40 Hz Coherence Decreases in REM Sleep and Ketamine Model of Psychosis. Front. Psychiatry 9, 766.

Hunt, M.J., Adams, N.E., Średniawa, W., Wójcik, D.K., Simon, A., Kasicki, S., and Whittington, M.A. (2019). The olfactory bulb is a source of high-frequency oscillations (130-180 Hz) associated with a subanesthetic dose of ketamine in rodents. Neuropsychopharmacology 44, 435–442.

Tort, A.B.L., Brankačk, J., and Draguhn, A. (2018). Respiration-Entrained Brain Rhythms Are Global but Often Overlooked. Trends Neurosci. 41, 186–197.

